# COLLAGE: COnsensus aLignment of muLtiplexing imAGEs

**DOI:** 10.1101/2024.07.15.603557

**Authors:** Asier Antoranz, Alexandre Arnould, Madhavi Dipak Andhari, Pouya Nazari, Gautam Shankar, Bart De Moor, Frederik De Smet, Francesca Maria Bosisio, Jon Pey

## Abstract

Multiplexed immunohistochemistry (mIHC) enables the high-dimensional single-cell interrogation of pathological tissue samples. mIHC is commonly based on the collection of high-resolution images from repeated staining cycles of the same tissue sample. Images of individual cycles typically consist of smaller tiles that need to be stitched into larger composite images, while images from serial rounds require alignment in a shared set of coordinates to enable pixel-perfect data integration. Current algorithms for stitching and registration require solving a single large puzzle consisting of billions of pixels making them computationally expensive but moreover forcing them to introduce errors to close the puzzle, which significantly impact the downstream results and the single-cell profiles. Here, we present the development and evaluation of COLLAGE (**CO**nsensus a**L**ignment of mu**L**tiplexing im**AGE**s), an innovative stitching and registration method that leverages on the complementarity of these two steps in a ‘divide and conquer’ approach: in contrast to other algorithms, COLLAGE breaks the process down into thousands of small puzzles, enabling extensive parallelisation and not forcing errors in its solution. Because COLLAGE also includes AlgnQC, a novel deep-learning-based evaluation metric of registration quality, the quality of the resulting image stacks is consistently maximised, while images with errors are flagged in an automated way. COLLAGE is available via www.disscovery.org.

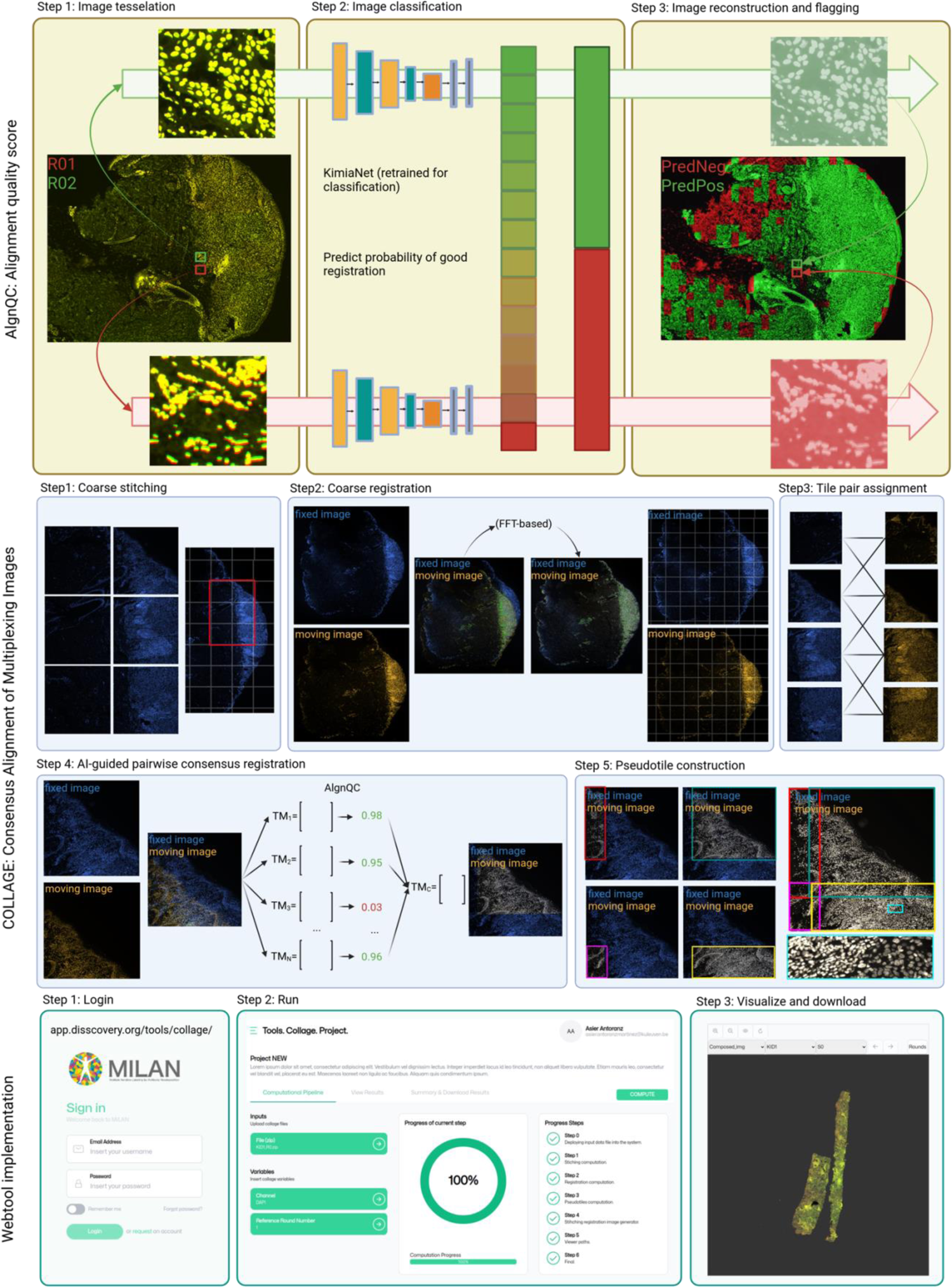

## Introduction

High-dimensional multiplexed immunohistochemistry (mIHC) is a powerful technique that enables the simultaneous visualization and analysis of multiple molecular targets within a single tissue sample, thereby providing a comprehensive understanding of the spatial distribution and interactions of cellular components within their native microenvironments (De Smet, F., et al., 2021), (Gerdes, M.J., et al., 2013), (Giesen, C., et al., 2014). This approach has been transformative in various research fields, including oncology, immunology, and neuroscience, by facilitating the study of complex cellular interactions, biomarker co-expression, and tissue heterogeneity (Angelo, M., et al., 2014), (Parra, E.R., et al., 2018), (Schulz, D., et al., 2018), (Naulaerts, S., et al., 2023), (Kruse, B., et al., 2023) and has been increasingly employed to unravel the complexity of biological systems and to identify new therapeutic targets, enhancing the potential for personalized medicine (Lin, J.R., et al., 2018), (Tsujikawa, T., et al., 2017), (Bodenmiller, B., 2016), (Antoranz, A., et al., 2022), (Lamarthee, B., et al., 2023).

Various approaches for mIHC have been developed to enable the simultaneous detection and analysis of multiple molecular targets in biological samples. These techniques can be broadly categorized into fluorescence-based (Cattoretti, G., et al., 2019), (Lin, J.R., et al., 2015), mass spectrometry-based (Giesen, C., et al., 2014), and nucleic acid-based methods (Chen, K.H., et al, 2015). Fluorescence-based approaches are among the most widespread techniques in mIHC-based imaging due to their ease of use, accessibility, and relatively low cost. These methods rely on the use of fluorophore-conjugated antibodies to target specific biomarkers, with the subsequent emission of distinct wavelengths of light that can be detected and analyzed using optical microscopy (Stack, E.C., et al., 2014), (Parra, E.R., et al., 2018). Although these methods offer high sensitivity and resolution, they are often limited by the spectral overlap between fluorophores, necessitating the use of advanced unmixing algorithms or cyclic staining and imaging protocols to resolve individual signals (Gut, G., et al., 2018), (Ruifrok, A.C., et al., 2001).

Each of these methods offers unique advantages and limitations. All-in-one acquisition techniques enable the rapid visualization of multiple biomolecules in a single imaging round (Stack, E.C., et al., 2014), which can be advantageous for preserving sample integrity and minimizing experimental time. However, the number of targets that can be simultaneously analyzed is often limited by the spectral overlap between fluorophores, reducing the multiplexing capabilities of this approach (Lin, J.R., et al., 2018). Cyclic acquisition techniques are the most widespread but, on the other hand, rely on iterative rounds of staining, imaging, and signal removal to increase the number of detectable targets. Examples of such techniques include cyclic immunofluorescence: MILAN (Cattoretti, G., et al., 2019, Bolognesi, M.M., et al., 2017), CycIF (Lin, J.R., et al., 2016, Gerdes, M.J., et al., 2013), CODEX (Goltsev, Y., et al., 2018, Black, S., et al., 2021), COMET (Lunaphore tm), multiplexed ion beam imaging (MIBI) (Angelo, M., et al., 2014), MIBI by time of flight (MIBI-TOF) (Keren, L., et al., 2019); and highly multiplexed RNA imaging methods such as multiplexed error-robust fluorescence in situ hybridization (MERFISH) and seqFISH (Schulz, D., et al., 2018). In these approaches, samples are stained and imaged multiple times, with each round targeting a different set of biomolecules.

By removing or inactivating the signal after each round, researchers can build a composite image of the tissue, revealing the spatial distribution of many targets. Although cyclic acquisition methods offer higher multiplexing capabilities, they can be more time-consuming than all-in-one acquisition techniques (Moffitt, J.R., et al., 2016), and require more complex workflows for analysis (Schapiro, D., et al., 2022). One of the most critical steps in these workflows is image registration where two or more images require pixel-perfect alignment to a common coordinate system (Zitová, B., & Flusser, J., 2003) ensuring that the spatial relationships between the different molecular targets are accurately maintained, allowing for a reliable projection of the quantification of these targets to the corresponding cells/locations. This step is crucial as it allows the extraction of the right molecular data in downstream analyses. This is typically obtained by subtracting the background obtained from baselines acquired in the first and last rounds (Potier, G., et al., 2022). As shown in Figure 01, if the images obtained during different cycles do not align, the signals obtained after background subtraction include signal that represents remains of the background rather than true signal. The larger registration errors become, the more erroneous also the molecular data extracted (typically in the form of MFI – Mean Fluorescence Intensity) for each cell in the different cycles becomes, including signals expressed by neighbouring cells.

**Figure 01.**
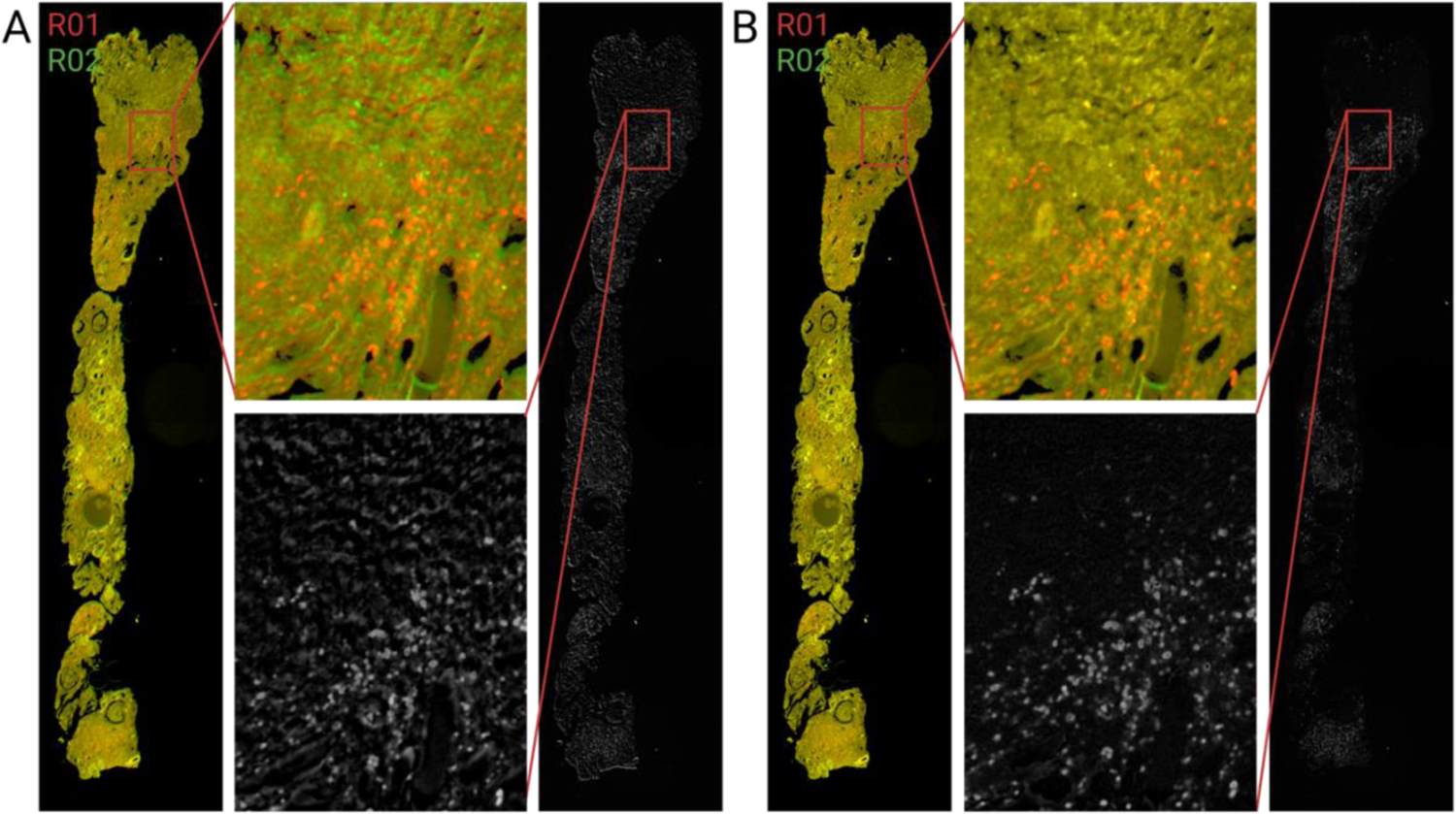
The importance of registration in the multiplexing image analysis workflow. **A)** incorrect background subtraction of a CD68 staining (red) acquired in the first cycle using an unstained baseline acquired in the next cycle. It can be seen how the shift between both images adds a lot of background signal to the processed image; B) the same but with correctly registered images. The processed image contains only signals corresponding to CD68 staining.

To overcome this challenge, researchers have developed various registration algorithms and software tools, some of which are specifically designed for multiplex imaging applications. These methods can be broadly classified into:

### Rigid registration

Involves the translation, rotation, and scaling of the reference and query images for the alignment. It can be further divided into:

### Manual registration

This is the simplest and most basic method, where images are registered manually by visually aligning the markers of interest (Maintz, J. B., & Viergever, M. A., 1998). Although manual registration is time-consuming and subject to human error, it can be useful for small datasets or when high precision is not required.

### Feature-based registration

This method involves identifying common features or landmarks in the images, such as blood vessels or nuclei, and using these features to align the images (Zitová, B., & Flusser, J., 2003). Popular algorithms such as SIFT (Scale-Invariant Feature Transform) (Lowe, D.G., et al., 2004) or SURF (Speeded Up Robust Features) (Bay, H., et al., 2008) are included in this category.

### Intensity-based registration

This method involves matching the intensities of the images to align them. It can be done using cross-correlation, mutual information, or other similarity measures (Pluim, J.P., et al., 2003).

### Elastic registration

This method involves using a deformation field to non-linearly warp the images to match each other (Sotiras, A., et al., 2013). It can be done using algorithms such as Thin Plate Splines (Bookstein, F.L., 1989) or Demons (Thirion, J.P., 1998).

### Hybrid methods

There are also several hybrid methods that combine different registration approaches to improve accuracy and robustness. For example, feature-based registration can be combined with elastic registration to correct for large distortions (Myronenko, A. & Song, X., 2010).

Slide scanners used in the acquisition of multiplexed images, do not capture the entire imaging surface at once, instead they acquire the imaging surface in a tessellated manner with tiles of a fixed size. Conventionally, tiles from the same scan are first stitched together and then the images from serial scans are registered. Newer methods aim to improve this process by integrating stitching and registration into a unified approach. Examples include Ashlar (Muhlich, J.L., et al., 2022) and AstroPath (Berry, S., et al., 2021). Despite their improvements, their approach of treating stitching and registration as a ‘single puzzle’ problem introduces certain limitations. While each pair of tiles can be accurately aligned/stitched, the collective alignment of the entire set sometimes leads to discrepancies. This may be caused by the miscalibration of the Z-axis, camera rotation, of overlapping sections of adjacent tiles belonging to different focal points (z-position slightly different) (Muhlich, J.L., et al., 2022). The core issue lies in the single-puzzle framework, where, by trying to stitch/register all the tiles in a single puzzle, they significantly constrain the solution space and when the entire puzzle does not fit together, the global optimum might be insufficient. This issue typically escalates with the size of the image; as the number of tiles increases, so does the potential for misalignment. Furthermore, these approaches have additional constraints (they only solve translation and require pre-processing to adjust scaling and rotation). To tackle these limitations, COLLAGE adopts a divide-and-conquer strategy by independently registering each pair of tiles. This splits the larger puzzle into potentially thousands of smaller puzzles which broadens the system’s degrees of freedom and thus providing a more adaptable, scalable, and error-resilient solution.

## Materials and Methods

### Data sets and availability

A set of well-curated datasets were considered for this study involving different cyclic multiplexing technologies, acquired in different laboratories, using different tissue types, at different resolutions. Dataset 1 (Lamarthee, B., et al., 2023) consists of a cohort of kidney transplant patients acquired with the MILAN (Cattoretti, G., et al., 2019, Bolognesi, M.M., et al., 2017) technology. The dataset is organised in 7 slides with 28 Core Needle Biopsies (CNBs). Dataset 2 consists of a multi cancer cohort acquired with the COMET technology (LunaPhore tm). The dataset is organised as one TMA with 9 cores (2xBRCA, 2xGBM, 2xHCC, 2xSKCM, 1xHealthyTonsil) and a whole resection of a healthy tonsil. Dataset 3 (Pozniak, J., et al., 2023) consists of a cohort of metastatic melanoma biopsies acquired with the CODEX (Goltsev, Y., et al., 2018) technology. The dataset is organised in 5 slides with 5 whole resection biopsies. Table 1 contains the complete set of details for each of the datasets.

**Table 1.** Datasets used to benchmark COLLAGE.

| Parameter | Dataset 1 | Dataset 2 | Dataset 3 |
| --- | --- | --- | --- |
| Acquisition technology | MILAN | COMET (LunaPhore) | CODEX (Akoya) |
| Number of cycles | 22 | 5 | 12 |
| Number of markers | 45 | 10 | 27 |
| Species | Homo Sapiens | Homo Sapiens | Homo Sapiens |
| Tissue type | Kidney transplant | Healthy Tonsil<br>PanCancer | Metastatic melanoma |
| Number of slides | 7 | 2 | 5 |
| Slide type | CNB | RB, TMA | RB |
| Laboratory | Laboratory 1 | Laboratory 1 | Laboratory 2 |
| Acquisition Instrument | Zeiss Axioscan Z1 | LunaPhore COMET | Keyence BZ-X800 |
| Resolution | 0.65 $\mu\text{m}/\text{px}$ | 0.22 $\mu\text{m}/\text{px}$ | 0.37 $\mu\text{m}/\text{px}$ |
| Tile size | 2040 x 2040 px | 2448 x 2048 px | 1920 x 1440 px |
| Overlapping adjacent tiles | ~10% | ~5.7% (horizontal)<br>~8.8% (vertical) | ~30% |

Prior to registration, all datasets have been pre-processed with the same pipeline. Image foreground was identified using a custom implementation of Coreograph, a foreground detection tool implemented in MCMICRO (Schapiro, D., et al., 2022) that was originally developed for TMAs but that we have expanded for any type of slide (TMA, CNB, RB). Tiles were corrected for background shading using BaSiC (Peng, T., et al., 2017).

### AIgnQC: A classification model for the evaluation of registration

In the context of evaluating registration performance in medical imaging, particularly in pathology, achieving a robust and generalizable score is crucial yet challenging. Several metrics have been suggested ranging from pixel-level error- and correlation-based metrics such as the median absolute error (MAE), mean square error (MSE), and Pearson correlation coefficient (R), to structure-based metrics such as the structural similarity index matrix (SSIM), mutual information coefficient (MI), or even hybrid metrics combining any of the previous statistics. These metrics, however, often fall short in capturing the variability inherent to tissue images as they have high case-to-case variability and are heavily correlated to the degree of error in the registration rather than providing dichotomic outputs. In this work, we leverage deep learning to surpass these limitations. Deep features, generated by a well-trained deep neural network, have consistently demonstrated superior performance over handcrafted features across various applications (Riasatian, A., et al., 2021, Jegou et al., 2011, Kumar et al., 2017).

In this study, we have trained a deep-learning classification model using the popular DenseNet-121 architecture (Huang, G., 2017). Here, transfer learning was applied by importing the weights from KimiaNet, a pretrained network for histopathology image representation (Riasatian, A., et al., 2021). The workflow used to train our model is represented in Figure 2A. Specifically, a subset of properly registered images was manually selected from all datasets (Table 1), and these were tessellated into tiles of 256px of lateral size. On top of visual evaluation, we calculated the cross correlation between each pair of registered tiles and removed those that were not located in the optimal location. In these datasets, the first round (R01) always represented the fixed image (reference/anchor), while the images belonging to the rest of the rounds represented the moving image (query). A correctly registered image with respect to the anchor was defined a ‘positive image’. For example, an image belonging to R02 correctly registered with R01 is defined a positive image. The combination of the anchor and the positive image made the positive composite (score = 1) for model training. Positive images were transformed (first translated and then rotated) using random values extracted from normal distributions: N_rotation_(0, 0.5), N_translation_(0, 10). Scaling was not applied since cyclic multiplexing scans always the same tissue which should not change in size. Transformed images were called ‘negative image’. The combination of the anchor and the negative image made the negative composite (score = 0) for model training. Following this approach 1,255,661 pairs of positive and negative composites were generated (824,471 for dataset 1, 249,5128 for dataset 2, and 181,672 for dataset 3). 100,000 tiles for each dataset were randomly selected for model training. The rest of tiles were used for testing. The model was then trained using the following parameters: batch_size = 8, optimized = Adam, learning_rate = 0.001, loss = ‘binary_crossentropy’, metrics = ‘accuracy’.

**Figure 2.**
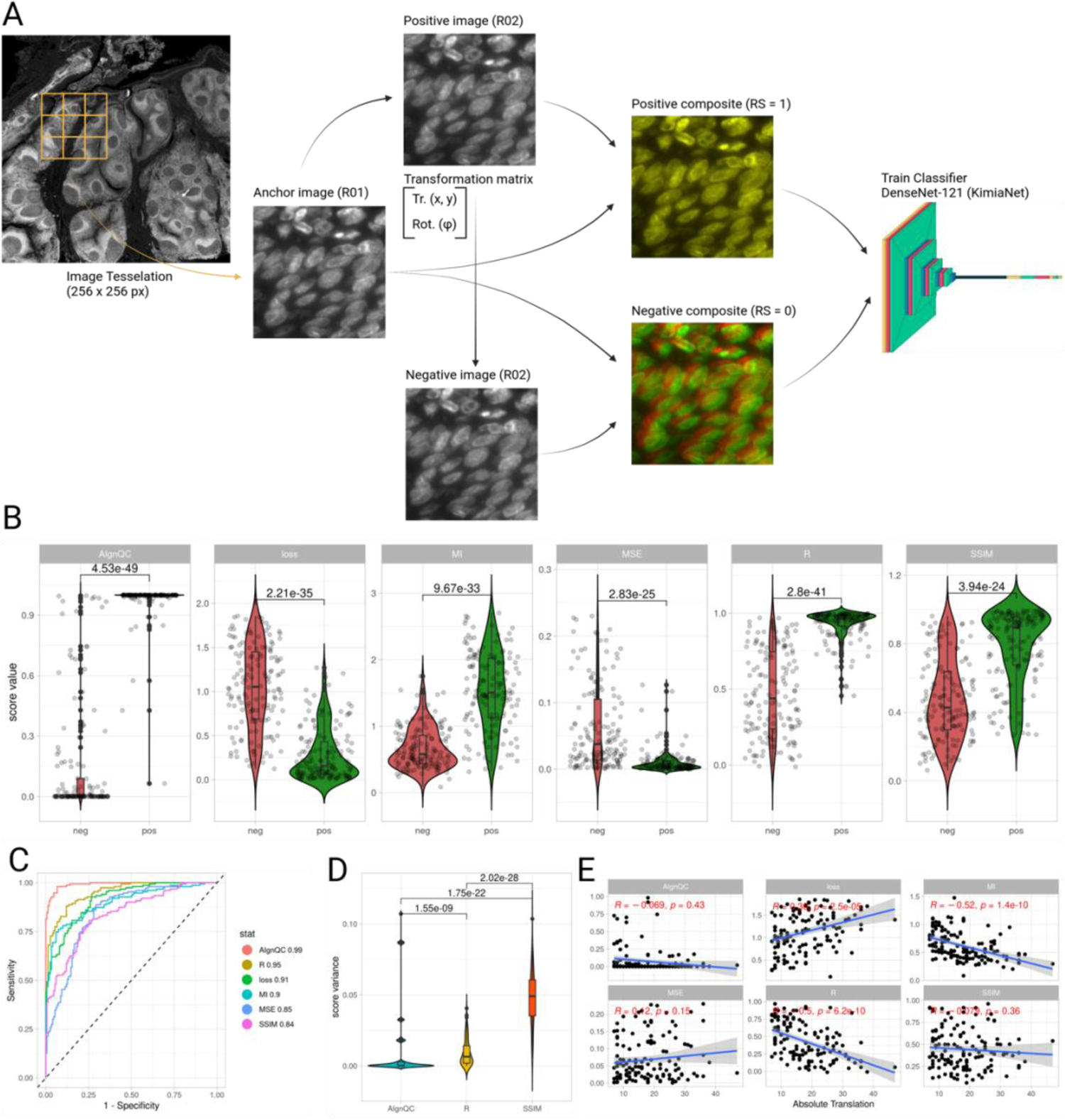
AIgnQC can classify good vs bad registration areas. A) Training scheme followed to train AlgnQC. Images properly registered were obtained and defined as ‘positive images’. These were transformed using a randomly generated transformation matrix (translation and rotation only) creating negative images. The composite of the reference image with the positive image represented the positive composites (RS = 1 for training). The (RS = 0 for training). The composite of the reference image with the negative image represented the negative composites (RS = 0 for training). A DenseNet-121 classification model was then trained using a pre-trained version with histopathological images (KimiaNet). B) The outcome of AlgnQC was compared to state-of-the-art scores used for the evaluation of registration showing that AlgnQC can reliably separate positively and negatively registered images. C) ROC curves comparing the AUC of AlgnQC with SOTA metrics. AlgnQC (AUC = 0.99) outperforms competing metrics. D) Comparison of variability of AlgnQC with SOTA metrics in positive images. The variability introduced by AlgnQC is significantly smaller than the one introduced by SOTA metrics making AlgnQC more robust. E) Correlation of SOTA metrics with the absolute introduced translation in the generation of negative images. AlgnQC lacks any correlation with the magnitude of the translation unlike other SOTA metrics.

After training, the model was evaluated in an independent set of samples from datasets 1-3. From each dataset, 50 randomly selected positive and negative composites were filtered to make meaningful comparisons between 6 different evaluation statistics including: MSE, SSIM, MI, R, loss, and our classification score. For the scores given by each of these statistics we calculated the AUC of a ROC curve separating positive vs negative cases as well as the p-value obtained from a Wilcoxon test comparing the scores for the positive vs the negative cases. To show the robustness and variability of each score, within the positive cases, we evaluated the variance of each score using bootstrapping analysis. To show the degree of dependence with respect to the registration error, within the negative cases, we evaluated the correlation between the score and the magnitude of translation introduced to generate the negative composites.

### COLLAGE: COnsensus aLignment of muLtiplexing imAGEs

The workflow followed by COLLAGE can be found in the graphical abstract. Specifically, COLLAGE follows these steps: 1) coarse stitching/registration for tile pair assignments, 2) Pairwise consensus registration, 3) Pseudotile construction.

#### Coarse stitching/registration for tile pair assignments

A key step in COLLAGE is to identify pairs of tiles from consecutive rounds with overlapping tissue. This is performed by coarsely stitching low-resolution reconstructions using the stage position of each tile and registering reference and query rounds. After the tiles have been stitched and registered, we assign all tile-pairs with overlapping tissue building a bipartite graph.

#### Pair-wise consensus registration

After all overlapping tiles have been identified, the next step consists of calculating a transformation matrix (translation, rotation, and scaling) for each pair of tiles with overlapping tissue to perform a rigid registration (this corresponds to every edge in the bipartite graph). Any rigid registration method involves spatially transforming the source image to align with the moving image and have 3 components: the geometric transformation, the similarity measure to be maximized, and an optimization technique. We have selected algorithms that vary in these three components.

– ImRegProc (Ri, Y., and Fijumoto, H., 2018): a robust image registration method using the Fast-Fourier-Transform (FFT) technique. The transformation matrix is estimated by first converting the images to the frequency domain and then inverting the transformation (iFFT) on the phase correlation between the two images. Finally, the Linear-Least-Squares (LLS) method is used to minimize the cost function and optimize the transformation matrix.
– ImReg (https://github.com/matejak/imreg_dft): an enhanced image registration tool that estimates the transformation matrix through two stages of phase correlation, excluding low spatial frequencies for effectiveness. Initially, it assesses angle and scale modifications using amplitude calculations of the Fourier spectrum followed by a log polar transformation and phase correlation. Subsequently, it conducts phase correlation on both the original and a 180° rotated version of the transformed image against the fixed image. The most accurate outcome between these comparisons determines the final adjustment angle and translation vector.
– Astroalign (Beroiz, M., et al., 2020): It is similar to feature-based image registration methods like SURF, SIFT or ORB where the correspondence of the located points is done using corner detection methods. Here instead, it focuses on finding and matching 3-point asterisms (triangles). Further, correspondence between the triangles is calculated using RANdom SAmpling Consensus (RANSAC), a method that proposes a transformation matrix that is in agreement with the majority of the matched features (triangles).
– ThunderReg (https://github.com/thunder-project/thunder-registration). Uses cross correlation to estimate an integer n-dimensional displacement between a query and a reference image.
– PyStackReg (https://pystackreg.readthedocs.io/en/latest/readme.html). It is an automatic subpixel registration algorithm that minimizes the mean square intensity difference between a reference and a moving image.
– AirLab (Sandkühler, R., et al., 2018). Open laboratory for image registration where the analytic gradients of the objective function are computed automatically.

Since the overlapping tissue between two tiles can sometimes be small, individual registration methods can often provide false positive results. Therefore, we apply our newly developed AlgnQC model to each method to identify these false positives (AlgnQC < 0.5). Among the methods with AlgnQC > 0.5, the transformation matrices are averaged in terms of translation, rotation, and scaling.

#### Pseudotile construction

finally, from each registered tile-pair, the overlapping region with the reference tile is cropped and merged with all the other query tile that overlap with the same reference tile. By doing this, we construct a pseudotile with ‘collaged’ pieces from several query tiles generating an equivalent to the reference image. This ‘divide-and-conquer’ procedure has the advantage that all the rounds are subject to the same final tile-stitching to reconstruct the whole image. Therefore, should any errors be forced, the same error will appear in all the rounds generating perfectly registered results.

COLLAGE has been implemented as a stand-alone tool and is available upon registration in the DISSCOvery set of tools (https://app.disscovery.org/login). Documentation on how to use the tool are available in the supplementary material.

## Results

### AIgnQC: a novel deep-learning model to assess the quality of registration

The evaluation of registration is often performed visually or using a series of statistics that require careful case-dependent adjustments during the evaluation process, making it challenging to exhaustively evaluate large datasets with tens of thousands of images. To streamline this process, we developed AlgnQC, a deep-learning-based quality score that evaluates the performance of the registration between multiplexing cycles. In this section we show the validity of AlgnQC. Specifically, we show that AlgnQC offers a better separation between correctly and incorrectly registered images than typically used statistics; then we show that AlgnQC is more robust and stable than other metrics; and finally, we show that AlgnQC does not depend on the magnitude of error in the registration.

First, we compared the classification score of AIgnQC with several statistics that are commonly used to evaluate the performance of registration methods. These metrics include error-based (MSE) and correlation-based (R) metrics as well as structure-based metrics (MI, SSIM), or hybrid metrics (loss = (1 – R) + (1 – SSIM)). An ideal score for image registration should have the following properties: a good separation between positive and negative cases, low variance within positive or within negative cases, lack of correlation between the score and the correctness of the registration.

To evaluate these, we first randomly sampled 50 positive and 50 negative image pairs. The scores associated to these images were compared between positive and negative cases using a Wilcoxon test and ROC analysis from which the AUC was extracted. Among all the evaluated metrics, AIgnQC had the lowest p-value (4.53e-49) and highest AUC (0.99) when compared with the rest of the evaluated metrics (Figure 2B and C). Second, the subset of positive cases was evaluated for variance. Among all the included scores, only R, SSIM, and AIgnQC have scores ranging up to 1 with 1 representing the perfect score. To demonstrate that AIgnQC has lower variance and is therefore more robust across different cases, we sampled 10 positive cases from the different datasets, and calculated the variance between the scores. This process was repeated 100 times, and we compared the set of variances between the different metrics. As shown in Figure 2D, AIgnQC had the lowest variance when compared to R (p-value = 1.55e-09) and SSIM (p-value = 1.75e-22). Third, we evaluated the subset of negative cases for correlation with the magnitude of translation introduced in the generation of the negative images. To that end, the magnitude of translation was calculated as the absolute value of the translations performed in x and y and this value was correlated with the statistic. Among all metrics, AIgnQC was the one with the lowest correlation (R = −0.069, p-value = 0.43) showing that is appropriate for classification tasks.

### The single puzzle paradigm often yields suboptimal results

Consider the toy example shown in Figure 3.A. This example involves 4 tiles (A, B, C, and D) arranged as a 2×2 grid. Adjacent tiles have approximately 10% of overlap on the edges to allow accurate stitching. We can calculate the transformation matrix to stitch each pair of tiles independently. The transformation matrix for a rigid 2D registration involving translation, rotation, and scaling can be represented as (http://web.cse.ohio-state.edu/~parent.1/classes/581/Lectures/5.2DtransformsAhandout.pdf):

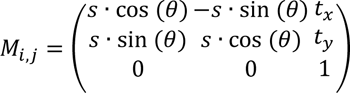

**Figure 3.**
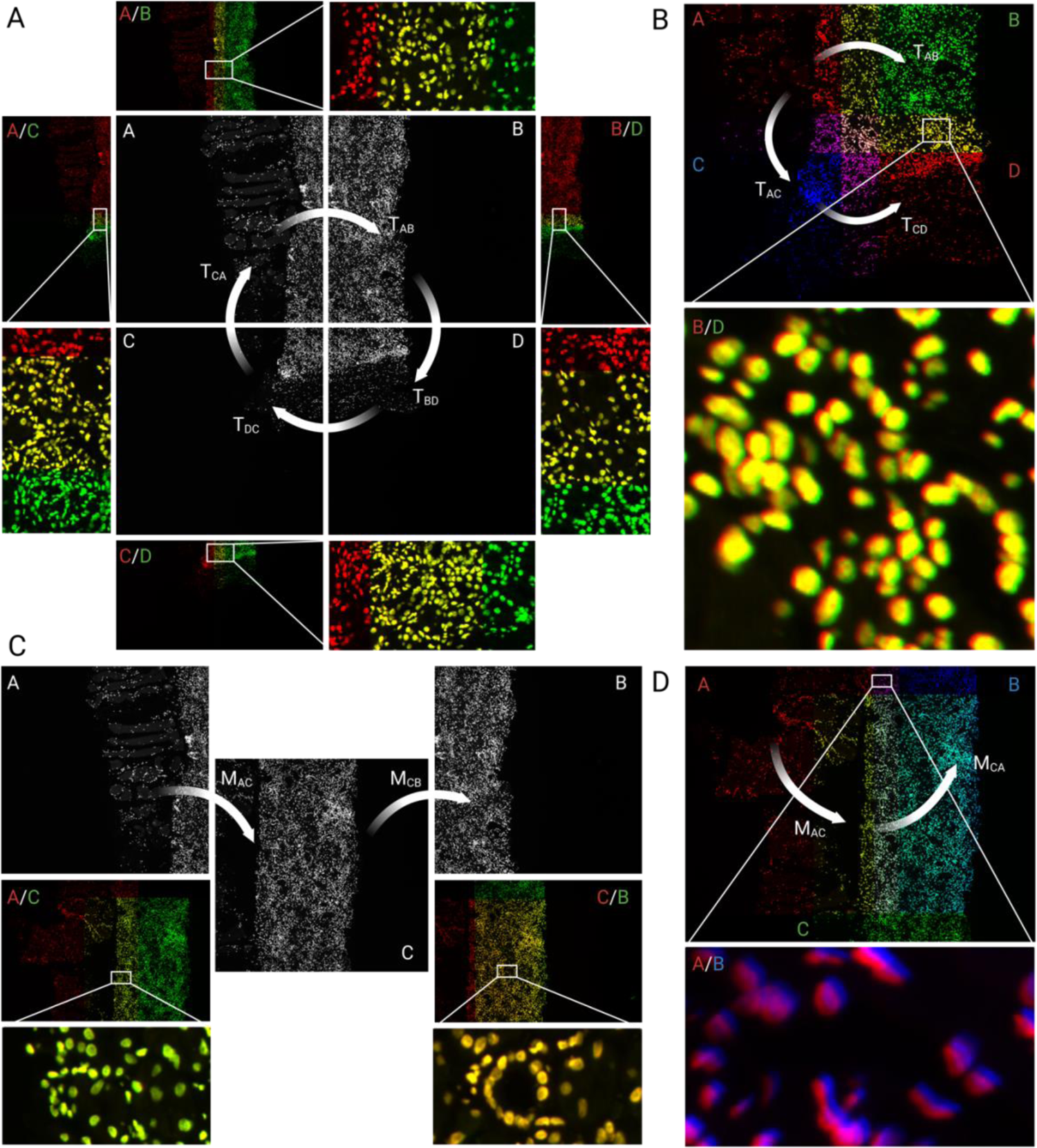
2D rigid registration: problem formulation and toy examples. A) First toy example involving 4 tiles from the same staining round arranged as a 2×2 grid. Adjacent tiles have approximately 10% of overlapping tissue for stitching. The zoomed-in areas show how the pairwise stitching of adjacent tiles leads to their pixel-perfect overlap. B) The single puzzle paradigm leads to suboptimal results in stitching as the cumulative translation vectors obtained from A to B and from A to C to D to B are different. C) Second toy example showing pixel-perfect overlapping for the pairwise registration of overlapping tiles. D) The single puzzle paradigm leads to suboptimal results in registration as the cumulative transformation matrix obtained from A to C to B and from A to B are different.

Where *s* is the scaling factor, θ is the rotation angle, and *t*_*x*_ and *t*_*y*_ the translations in the x and y axes, respectively. To calculate the cumulative effect of applying these transformations in a loop (A → B → D → C → A), the transformation matrices need to be orderly multiplied as follows:

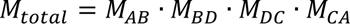

Where *M*_*AB*_ is the transformation matrix aligning tiles A and B. In the case of stitching, we are aligning tiles of the same round and therefore we can assume that s = 1 and θ = 0 so we can simplify the matrixial representation to a vectorial representation:

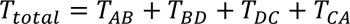

Where *T*_*AB*_ = (*T*_*x*_, *T*_*y*_) is the translation vector aligning tiles A and B. If the system of alignments represents a ‘closing puzzle’, *i.e.,* the system of transformations brings us back to the starting point without error, then the sum of translations applied sequentially around the loop of tiles results in a zero total translation (*T*_*total*_ = 0). For matrixial representation, the equivalent would be the identity matrix (*M*_*total*_ = *I*).

As shown in Table 2.A, the loop does not close. This is illustratively represented in Figure 3.B where the tiles were stitched following two independent paths. On the one hand, we calculated the translation from A to B. On the other hand, we calculated the translations from A to C and from C to D. B and D have around 10% of overlapping tissue. If the puzzle closes, B and D should align perfectly without the need to further translate the images. As shown in Figure 3.B, since B and D do not align, the puzzle doesn’t close.

**Table 2.** Toy example results. A) Translations for toy-example 1. B) Transformations for toy example 2.

| Translation | T<sub>x</sub> | T<sub>y</sub> |  | Transformation | T<sub>x</sub> | T<sub>y</sub> | θ |
| --- | --- | --- | --- | --- | --- | --- | --- |
| T<sub>AB</sub> | +1829 | -1 |  | M<sub>AC</sub> | +1335 | +239 | -0.27 |
| T<sub>BD</sub> | +2 | +1827 |  | M<sub>CB</sub> | +495 | -237 | +0.26 |
| T<sub>DC</sub> | -1828 | +1 |  | M<sub>BA</sub> | -1829 | +1 | 0 |
| $T_{CA}$ | -1 | -1826 | | | | | |
| $T_{total}$ | 2 | 1 | | $M_{total}$ | +1 | +3 | -0.01 |

Consider now the toy example of Figure 3.C. In this case we have three tiles belonging to two different rounds. A and B are the same tiles from the first toy example, while tile C corresponds to a tile of the next round. Both rounds have been scanned at the same resolution. Therefore, we can simplify the affine transformation matrix to consider only translation and rotation (s = 1). In this case, the loop (A → C → B → A) could be represented as follows:

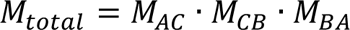

Similarly as before, if the system of alignments represents a ‘closing puzzle’, the sequential product of transformations around the loop of tiles results in the identity matrix bringing us back to the starting point without error, that is, *M*_*total*_ = *I*.

As shown in Table 2.B, the loop does not close. This is illustratively represented in Figure 3.D where the tiles were registered from A to C and from C to B. A and B belong to the same round and the translation from A to B has already been calculated in the previous toy example. A and B have around 10% of overlapping tissue. If the puzzle closes, A and B should align perfectly without the need to further translate the images. As shown in Figure 3.D, since A and B do not align, the puzzle doesn’t close.

These two toy examples highlight the limitations of the single-puzzle paradigm. These toy examples consider only small sets of tiles, while whole slide images often involve hundreds of tiles. Errors picked up at any stage accumulate by agglomerating the system which results in the reduction of its degrees of freedom. Newer methodologies like Ashlar (Muhlich, J.L., et al., 2022) and AstroPath (Berry, S., et al., 2021) do not follow this agglomerative approach and minimize the global error but still follow the single-puzzle paradigm. Therefore, COLLAGE follows a divide and conquer strategy by breaking down the single-puzzle paradigm into multiple systems and builds the images by collage-ing image patches that have been registered/stitched in a pairwise manner.

### COLLAGE outperforms state-of-the-art methods in registration

After a proper metric for registration has been defined, we compared the performance of COLLAGE with state-of-the-art open-source rigid registration methods including Ashlar (Muhlich, J.L., et al., 2022), AstroAlign (Beroiz, M., et al., 2020), ImReg (https://github.com/matejak/imreg_dft), PyStackReg (https://pystackreg.readthedocs.io/en/latest/readme.html), and ThunderReg (https://github.com/thunder-project/thunder-registration). The comparison was performed in the three different datasets as described above. In each dataset, the percentage of patches with an AlgnQC score above 0.1 was calculated. Only patches with at least 10% of foreground tissue were considered for the comparison. For the COMET and CODEX datasets (Figure 4A, and D (mid)), only ASHLAR and COLLAGE were considered. For the MILAN dataset, all registration methods were used. The COMET dataset included a WSI of a tonsil and a TMA with different cancer types. COLLAGE outperformed ASHLAR (Wilcoxon p-adj = 0.0134). While both methods provided very good results (mean score: ASHLAR = 0.996, COLLAGE = 0.999), ASHLAR still included certain regions where the tissue was not correctly registered (Figure 4A). In the CODEX dataset both methods performed comparably (p-adj = 0.0744) with COLLAGE providing slightly higher scores (mean score: ASHLAR = 0.994, COLLAGE = 0.996). The MILAN dataset involves CNBs which often is the most challenging dataset for registration due to their large eccentricity of the tissue samples (high length/width ratio) which causes that a lot of tiles pairs have very little information to perform the stitching/registration. In this case, COLLAGE (mean score = 0.983) outperformed all the other methods: ASHLAR (mean score = 0.317, p.adj = 3.19e-120), AstroAlign (mean score = 0.662, p.adj = 5.88e-51), ImReg (mean score = 0.876, p.adj = 1.27e-34), PyStackReg (mean score = 0.747, p.adj = 5.44e-38), ThunderReg (mean score = 0.271, p.adj = 7.34e-131).

**Figure 4.**
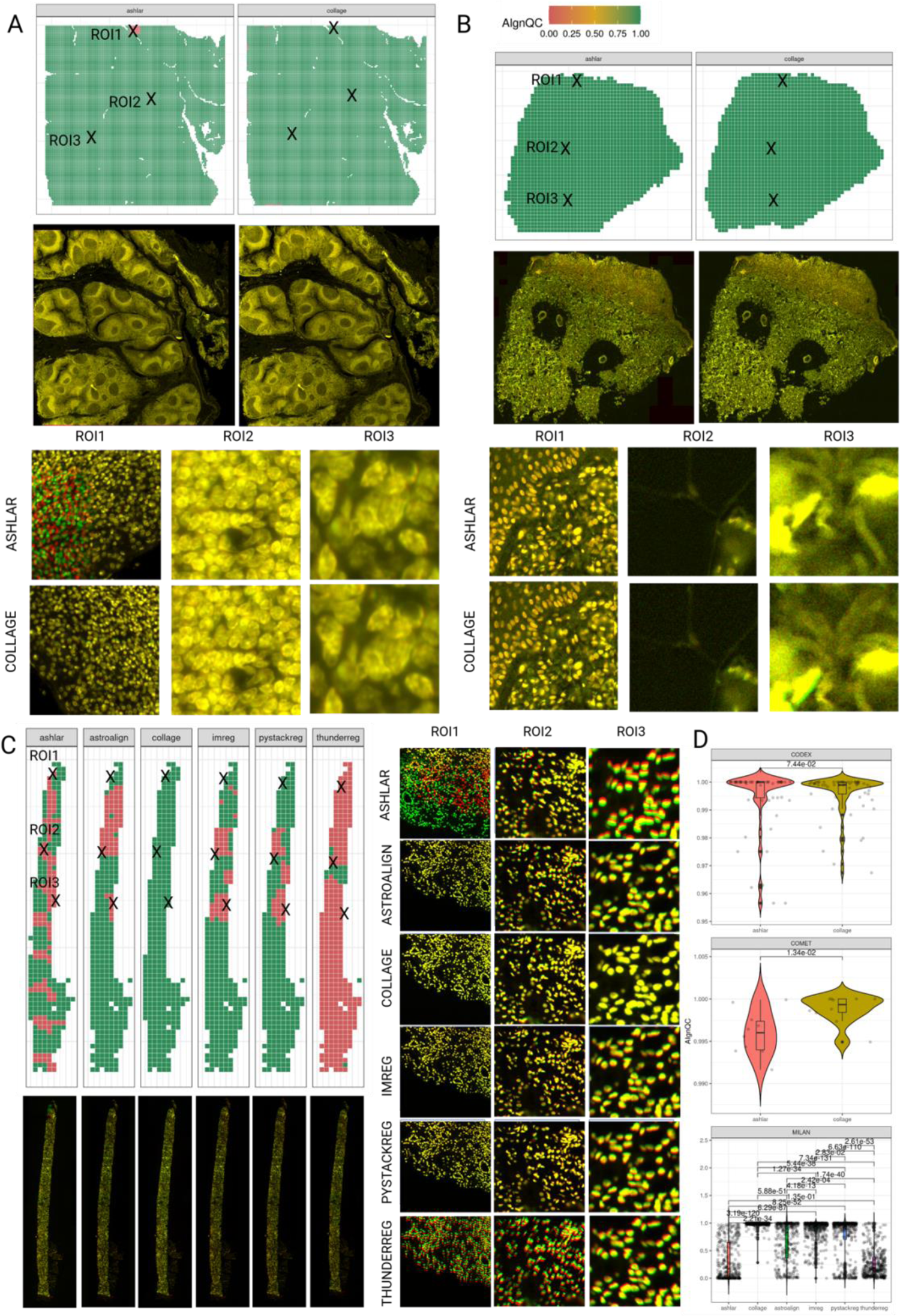
COLLAGE outperforms SOTA rigid registration methods. A) Comparison between Ashlar and COLLAGE in Lunaphore Comet data, B) Comparison between Ashlar and COLLAGE in Akoya Codex data, C) Comparison between COLLAGE and Ashlar, AstroAlign, ImReg, PyStackReg, and ThunderReg. D) Boxplots showing the differences in AlgnQC score between the different methods in the datasets included in the study.

## Conclusions

In this study we have shown that capturing the performance of registration methods with traditional metrics is very case-specific where a cut-off that might work for one case won’t work for another case making the identification of poorly registered areas difficult to automate. Here, we have shown how deep learning can overcome this limitation by introducing a novel classification score (AlgnQC) which is able to flag poorly registered areas with high accuracy for a very heterogeneous set of cohorts.

We also show how COLLAGE outperforms other methods in different scenarios. While Ashlar performs also very well for whole resection biopsies in whole image scans (usually denser tissue with more information), it still includes certain regions where the tissue is not correctly registered. We hypothesize that the minimal spanning tree construction used by Ashlar leaves these regions disjoint from the main connected component of the graph leading to these poorly registered areas. Additionally, it seems to struggle with less information dense tissues such as CNBs.

COLLAGE introduces a change of paradigm in the solution of stitching/registrations breaking out from the traditional ‘single-puzzle’ paradigm. Additionally, COLLAGE benefits from a consensus-based approach. We show how individual registration methods often return different transformation matrices and how a consensus implementation reduces the number of potential false positives.

The methods used here are for rigid registration (translation, rotation, and scaling). The authors recognize the limitations associated with rigid registration when physical deformation of the tissue occurs. In these cases, a Geometric distortion is still needed as a final correction. This issue affects all registration methods but splitting the issue into several problems as done in COLLAGE limits the errors to the areas affected.

Additionally, we provide COLLAGE as a webtool that can be used by non-experts in the field with an attractive user interface. A comprehensive tutorial is included in supplementary data 1.

## Acknowledgements

AA is supported by an FWO PhD fellowship fundamental research (12ATN24N). This work is supported by KULeuven (C3/ grant), het Vlaamse Veerkracht programma (VLAIO), “We also wish to acknowledge the “Opening the Future” campaign of KU Leuven for financial support.

## References

Angelo, M., et al. (2014). Multiplexed ion beam imaging of human breast tumors. Nature Medicine, 20(4), 436–442.

Antoranz, A., Van Herck, Y., Bolognesi, M.M., Lynch, S.M., Rahman, A., Gallagher, W.M., Boecxstaens, V., Marine, J.C., Cattoretti, G., van den Oord, J.J. and De Smet, F., 2022. Mapping the immune landscape in metastatic melanoma reveals localized cell–cell interactions that predict immunotherapy response. Cancer Research, 82(18), pp.3275–3290.

Avants, B. B., et al. (2011). A reproducible evaluation of ANTs similarity metric performance in brain image registration. NeuroImage, 54(3), 2033–2044.

Bay, H., Ess, A., Tuytelaars, T., & Van Gool, L. (2008). Speeded-up robust features (SURF). Computer Vision and Image Understanding, 110(3), 346–359.

Beroiz, M., Cabral, J.B. and Sanchez, B., 2020. Astroalign: A Python module for astronomical image registration. Astronomy and Computing, 32, p.100384.

Berry, S., Giraldo, N.A., Green, B.F., Cottrell, T.R., Stein, J.E., Engle, E.L., Xu, H., Ogurtsova, A., Roberts, C., Wang, D. and Nguyen, P., 2021. Analysis of multispectral imaging with the AstroPath platform informs efficacy of PD-1 blockade. Science, 372(6547), p.eaba2609.

Black, S., et al. (2021). CODEX multiplexed tissue imaging with DNA-conjugated antibodies. Nat. Protoc. 16, 3802–3835.

Bodenmiller, B. (2016). Multiplexed epitope-based tissue imaging for discovery and healthcare applications. Cell Systems, 2(4), 225–238.

Bolognesi, M.M., Manzoni, M., Scalia, C.R., Zannella, S., Bosisio, F.M., Faretta, M. and Cattoretti, G., 2017. Multiplex staining by sequential immunostaining and antibody removal on routine tissue sections. Journal of Histochemistry & Cytochemistry, 65(8), pp.431–444.

Bookstein, F. L. (1989). Principal warps: thin-plate splines and the decomposition of deformations. IEEE Transactions on Pattern Analysis and Machine Intelligence, 11(6), 567–585.

Cattoretti, G., Bosisio, F.M., Marcelis, L. and Bolognesi, M.M., 2019. Multiple iterative labeling by antibody neodeposition (MILAN).

Chen, K. H., et al. (2015). RNA imaging. Spatially resolved, highly multiplexed RNA profiling in single cells. Science, 348(6233), aaa6090.

De Smet, F., Martinez, A.A. and Bosisio, F.M., 2021. Next-generation pathology by multiplexed immunohistochemistry. Trends in Biochemical Sciences, 46(1), pp.80–82.

Gerdes, M. J., et al. (2013). Highly multiplexed single-cell analysis of formalin-fixed, paraffin-embedded cancer tissue. Proceedings of the National Academy of Sciences, 110(29), 11982–11987.

Giesen, C., et al. (2014). Highly multiplexed imaging of tumor tissues with subcellular resolution by mass cytometry. Nature Methods, 11(4), 417–422.

Goltsev, Y., Samusik, N., Kennedy-Darling, J., Bhate, S., Hale, M., Vazquez, G., Black, S. and Nolan, G.P., 2018. Deep profiling of mouse splenic architecture with CODEX multiplexed imaging. Cell, 174(4), pp.968–981.

Gut, G., et al. (2018). Multiplexed imaging of high-density libraries by sequential barcoded fluorescence in situ hybridization. Proceedings of the National Academy of Sciences, 115(42), E9889–E9898.

Huang, G., Liu, Z., Van Der Maaten, L. and Weinberger, K.Q., 2017. Densely connected convolutional networks. In Proceedings of the IEEE conference on computer vision and pattern recognition (pp. 4700–4708).

Jégou, H., Perronnin, F., Douze, M., Sánchez, J., Pérez, P. and Schmid, C., 2011. Aggregating local image descriptors into compact codes. IEEE transactions on pattern analysis and machine intelligence, 34(9), pp.1704–1716.

Keren, L., Bosse, M., Thompson, S., Risom, T., Vijayaragavan, K., McCaffrey, E., Marquez, D., Angoshtari, R., Greenwald, N.F., Fienberg, H. and Wang, J., 2019. MIBI-TOF: A multiplexed imaging platform relates cellular phenotypes and tissue structure. Science advances, 5(10), p.eaax5851.

Kruse, B., Buzzai, A.C., Shridhar, N., Braun, A.D., Gellert, S., Knauth, K., Pozniak, J., Peters, J., Dittmann, P., Mengoni, M. and van der Sluis, T.C., 2023. CD4+ T cell-induced inflammatory cell death controls immune-evasive tumours. Nature, pp.1–8.

Kumar, M.D., Babaie, M., Zhu, S., Kalra, S. and Tizhoosh, H.R., 2017, November. A comparative study of CNN, BoVW and LBP for classification of histopathological images. In 2017 IEEE symposium series on computational intelligence (SSCI) (pp. 1-7). IEEE.

Lamarthée, B., Callemeyn, J., Van Herck, Y., Antoranz, A., Anglicheau, D., Boada, P., Becker, J.U., Debyser, T., De Smet, F., De Vusser, K. and Eloudzeri, M., 2023. Transcriptional and spatial profiling of the kidney allograft unravels a central role for FcyRIII+ innate immune cells in rejection. Nature Communications, 14(1), p.4359.

Lin, J. R., Fallahi-Sichani, M., & Sorger, P. K. (2015). Highly multiplexed imaging of single cells using a high-throughput cyclic immunofluorescence method. Nature Communications, 6(1), 1–11.

Lin, J.R., et al. (2016). Cyclic Immunofluorescence (CycIF), A Highly Multiplexed Method for Single-cell Imaging. Curr. Protoc.Chem. Biol. 8, 251–264.

Lin, J. R., et al. (2018). Highly multiplexed immunofluorescence imaging of human tissues and tumors using t-CyCIF and conventional optical microscopes. eLife, 7, e31657.

Lowe, D. G. (2004). Distinctive image features from scale-invariant keypoints. International Journal of Computer Vision, 60(2), 91–110.

Maintz, J. B., & Viergever, M. A. (1998). A survey of medical image registration. Medical Image Analysis, 2(1), 1–36.

Moffitt, J. R., et al. (2016). High-throughput single-cell gene-expression profiling with multiplexed error-robust fluorescence in situ hybridization. Proceedings of the National Academy of Sciences, 113(39), 11046–11051.

Muhlich, J.L., Chen, Y.A., Yapp, C., Russell, D., Santagata, S. and Sorger, P.K., 2022. Stitching and registering highly multiplexed whole-slide images of tissues and tumors using ASHLAR. Bioinformatics, 38(19), pp.4613–4621.

Myronenko, A. and Song, X., 2010. Point set registration: Coherent point drift. IEEE transactions on pattern analysis and machine intelligence, 32(12), pp.2262–2275.

Naulaerts, S., Datsi, A., Borras, D.M., Antoranz Martinez, A., Messiaen, J., Vanmeerbeek, I., Sprooten, J., Laureano, R.S., Govaerts, J., Panovska, D. and Derweduwe, M., 2023. Multiomics and spatial mapping characterizes human CD8+ T cell states in cancer. Science Translational Medicine, 15(691), p.eadd1016.

Parra, E. R., et al. (2018). Validation of multiplex immunofluorescence panels using multispectral microscopy for immune-profiling of formalin-fixed and paraffin-embedded human tumor tissues. Scientific Reports, 8(1), 1–11.

Peng, T., et al. (2017). A BaSiC tool for background and shading correction of optical microscopy images. Nature communications, 8(1), p.14836.

Pluim, J. P., Maintz, J. B., & Viergever, M. A. (2003). Mutual-information-based registration of medical images: a survey. IEEE Transactions on Medical Imaging, 22(8), 986–1004.

Potier, G., Doméné, A. and Paul-Gilloteaux, P., 2022. A flexible open-source processing workflow for multiplexed fluorescence imaging based on cycles. F1000Research, 11.

Pozniak, J., Pedri, D., Landeloos, E., Van Herck, Y., Antoranz, A., Vanwynsberghe, L., Nowosad, A., Roda, N., Makhzami, S., Bervoets, G. and Maciel, L.F., 2024. A TCF4-dependent gene regulatory network confers resistance to immunotherapy in melanoma. Cell, 187(1), pp.166–183.

Ri, Y. and Fujimoto, H., 2018, March. Drift-free motion estimation from video images using phase correlation and linear optimization. In 2018 IEEE 15th International Workshop on Advanced Motion Control (AMC) (pp. 295-300). IEEE.

Riasatian, A., Babaie, M., Maleki, D., Kalra, S., Valipour, M., Hemati, S., Zaveri, M., Safarpoor, A., Shafiei, S., Afshari, M. and Rasoolijaberi, M., 2021. Fine-tuning and training of densenet for histopathology image representation using tcga diagnostic slides. Medical Image Analysis, 70, p.102032.

Ruifrok, A. C., & Johnston, D. A. (2001). Quantification of histochemical staining by color deconvolution. Analytical and Quantitative Cytology and Histology, 23(4), 291–299.

Sandkühler, R., Jud, C., Andermatt, S. and Cattin, P.C., 2018. AirLab: autograd image registration laboratory. arXiv preprint arXiv:1806.09907.

Schapiro, D., et al., 2022. MCMICRO: a scalable, modular image-processing pipeline for multiplexed tissue imaging. Nature methods, 19(3), pp.311–315.

Schulz, D., et al. (2018). Simultaneous multiplexed imaging of mRNA and proteins with subcellular resolution in breast cancer tissue samples by mass cytometry. Cell Systems, 6(1), 25–36.

Sotiras, A., et al. (2013). Deformable medical image registration: A survey. IEEE Transactions on Medical Imaging, 32(7), 1153–1190.

Stack, E. C., Wang, C., Roman, K. A., & Hoyt, C. C. (2014). Multiplexed immunohistochemistry, imaging, and quantitation: A review, with an assessment of Tyramide signal amplification, multispectral imaging and multiplex analysis. Methods, 70(1), 46–58.

Thirion, J. P. (1998). Image matching as a diffusion process: an analogy with Maxwell’s demons. Medical Image Analysis, 2(3), 243–260.

Tsujikawa, T., et al. (2017). Quantitative multiplex immunohistochemistry reveals myeloid-inflamed tumor-immune complexity associated with poor prognosis. Cell Reports, 19(1), 203–217.

Zitová, B., & Flusser, J. (2003). Image registration methods: a survey. Image and Vision Computing, 21(11), 977–1000.

