## supplementary material for "COLLAGE: COnsensus aLignment of muLtiplexing imAGEs"

### COLLAGE USER MANUAL

#### Registration & Login

Surf to https://app.disscovery.org/. There, they will be greeted with a welcome screen where you can use your credentials to login. If you do not have an account yet, you can request a new one with the "request an account" button.

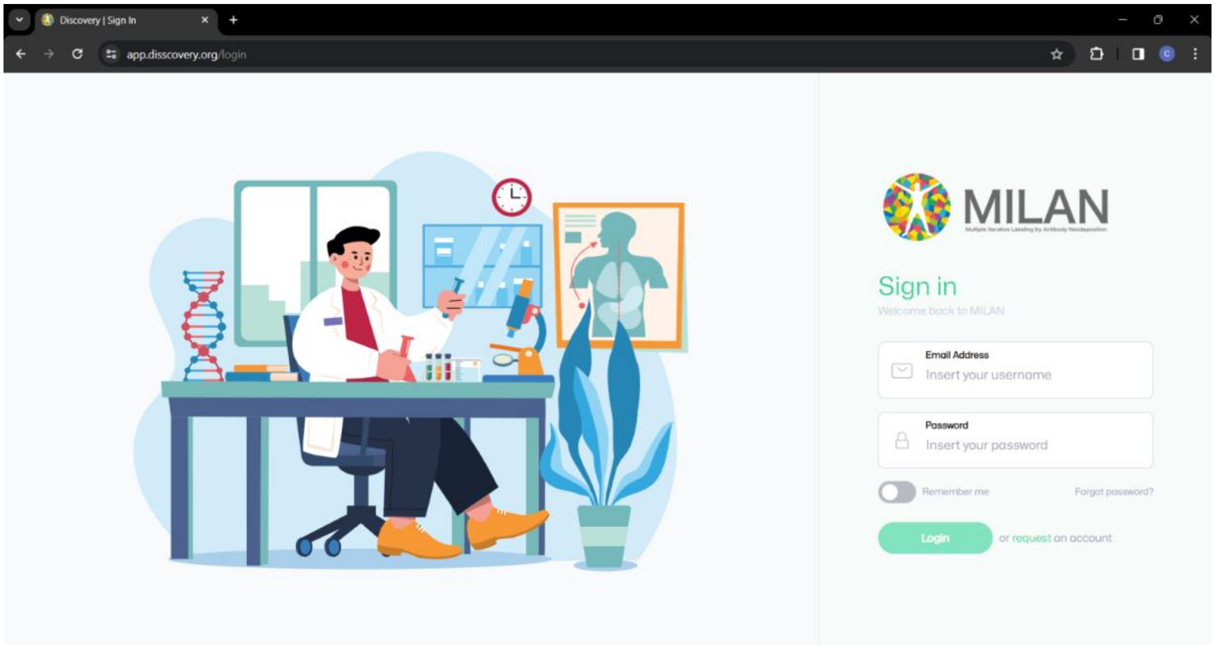


After login in, go to "Disscovery tools"

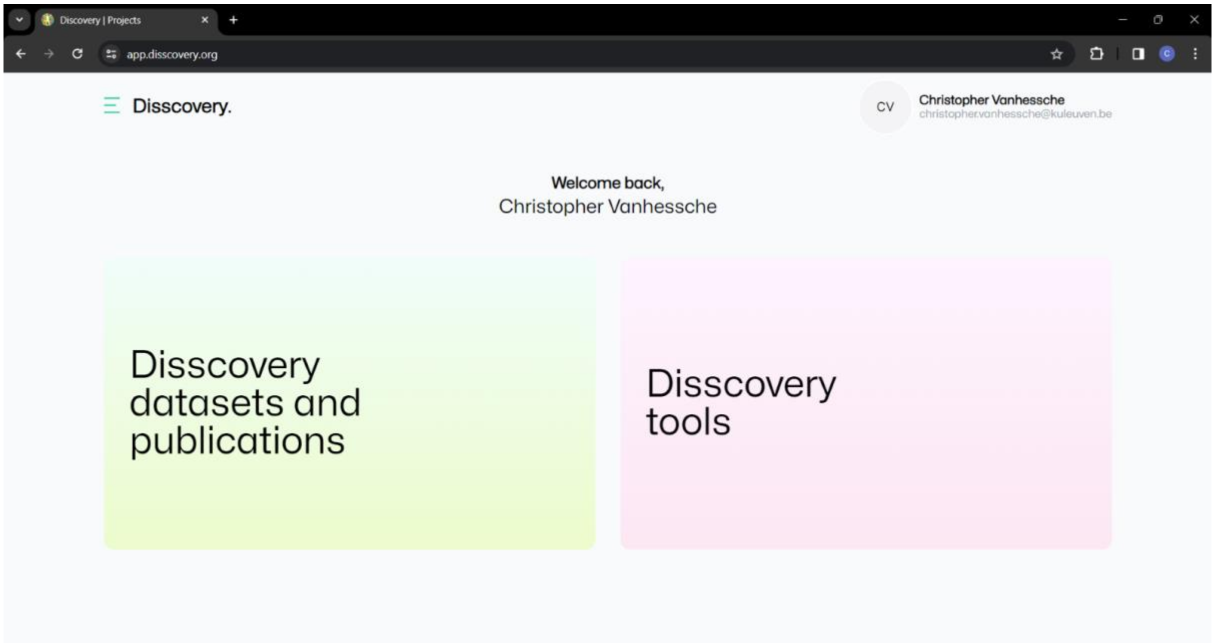

Click on “COLLAGE”.

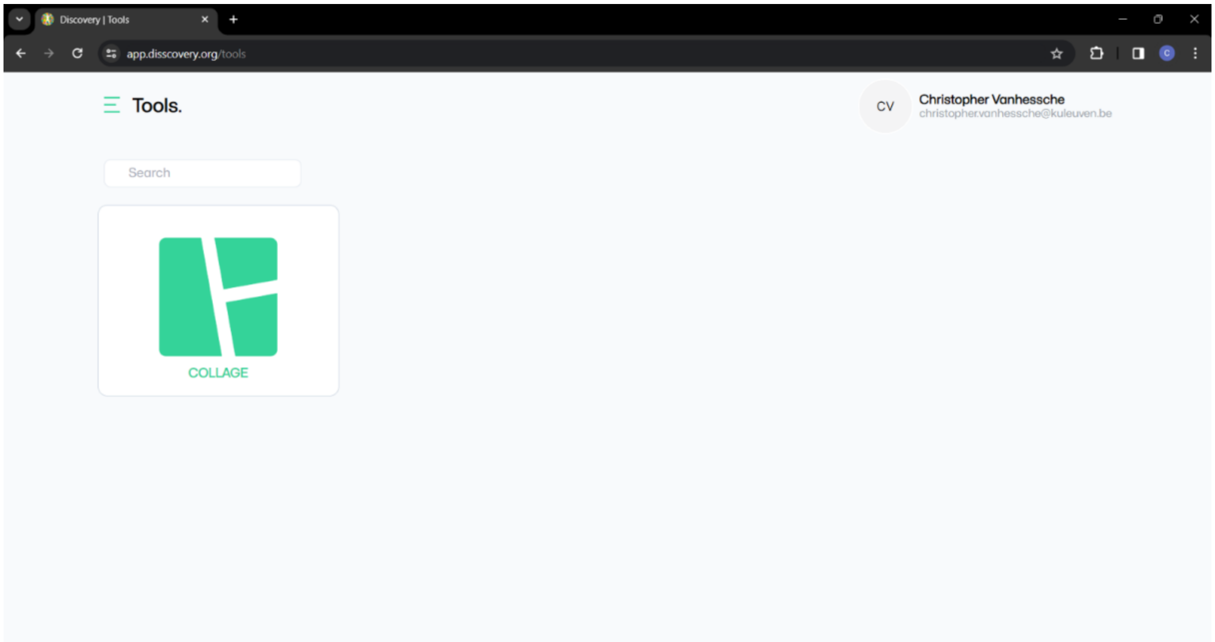


#### Project creation and setup

This will show your personal workspace. Here you can create a new project by clicking the “Add New Project” button.

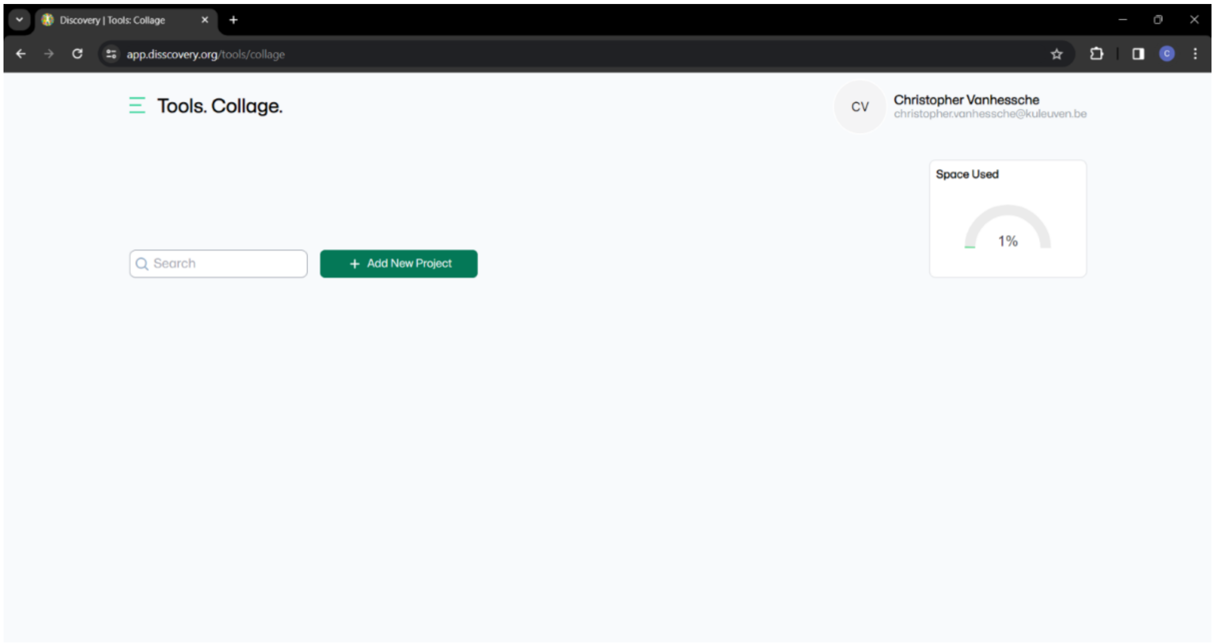


Here, you will be asked to name your project and briefly describe it. The Project Type should be set to “Disscovery” to benefit from Collage’s cloud computing options.

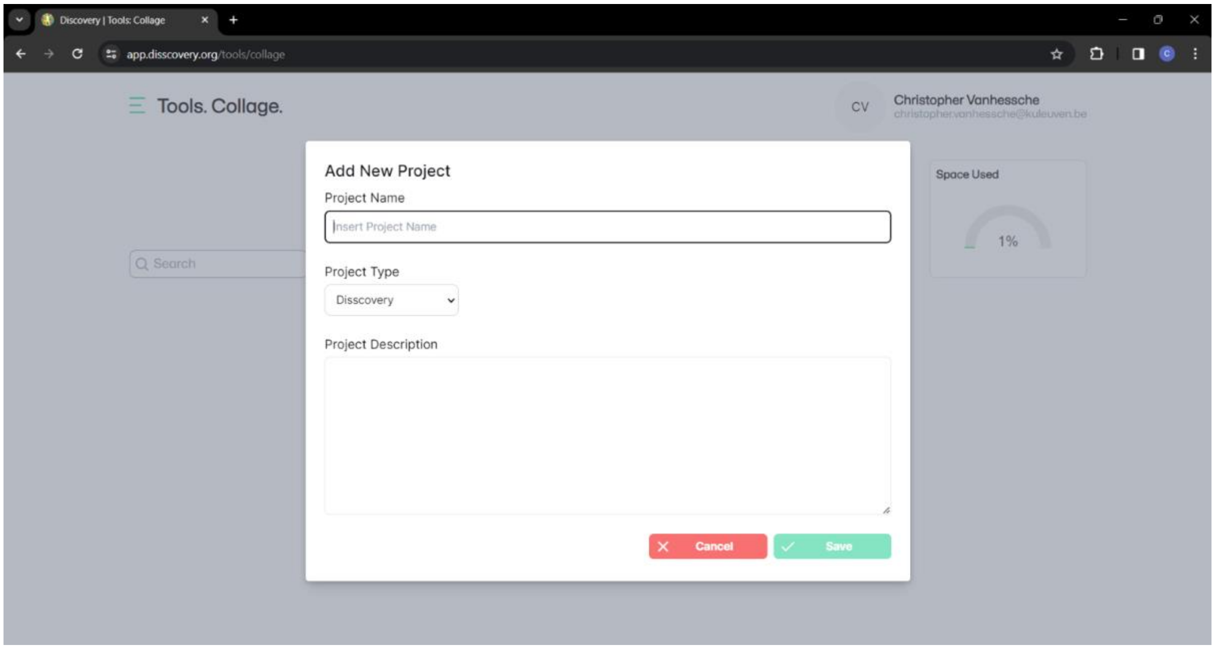


Setting the Project Type parameter to “Milan tool” assumes that the project-related data will be loaded from a local path which requires a more complex setup. If that is of interest to you, please contact.

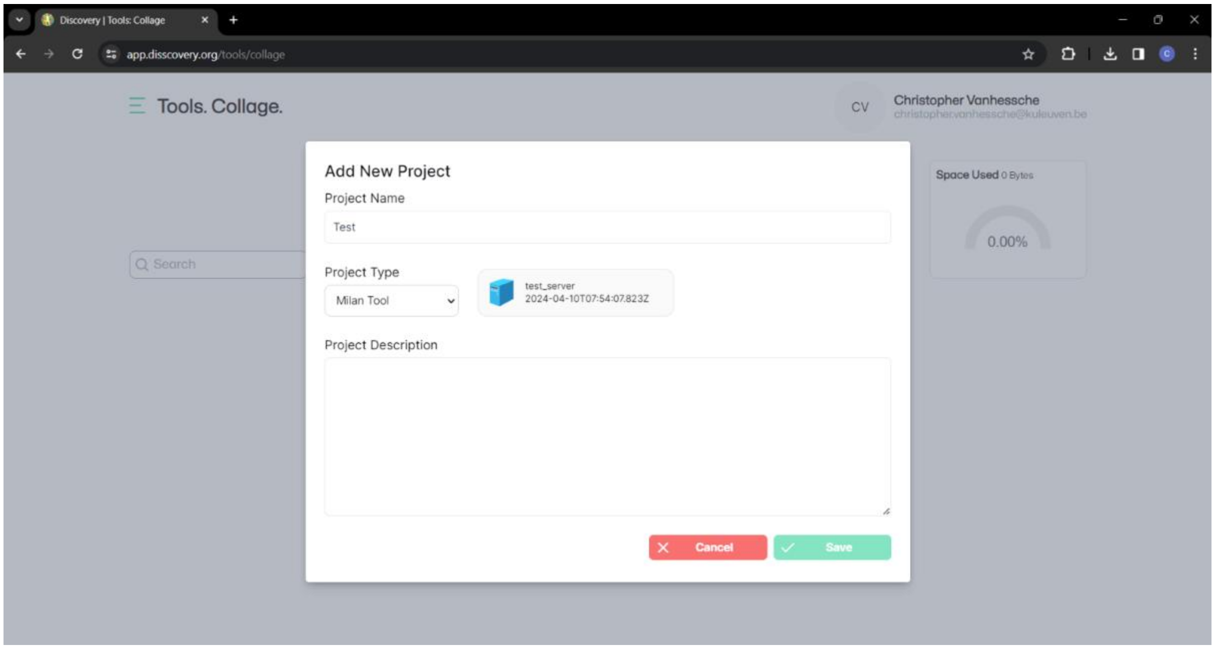


The created project can now be accessed via “View Project”.

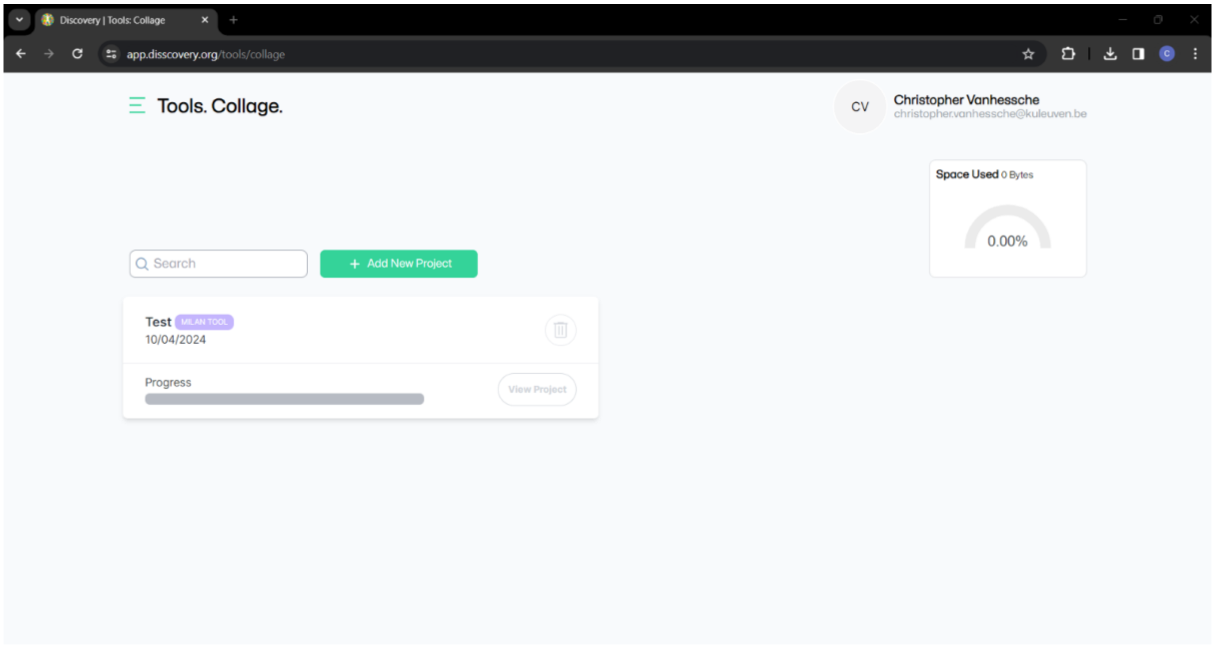


By accessing the project view you can see a complete overview of your project. This includes the computation steps, visualization and downloads options, among others. Note that 'View Results' and 'Summary & Download Results' are not available until the results are computed.

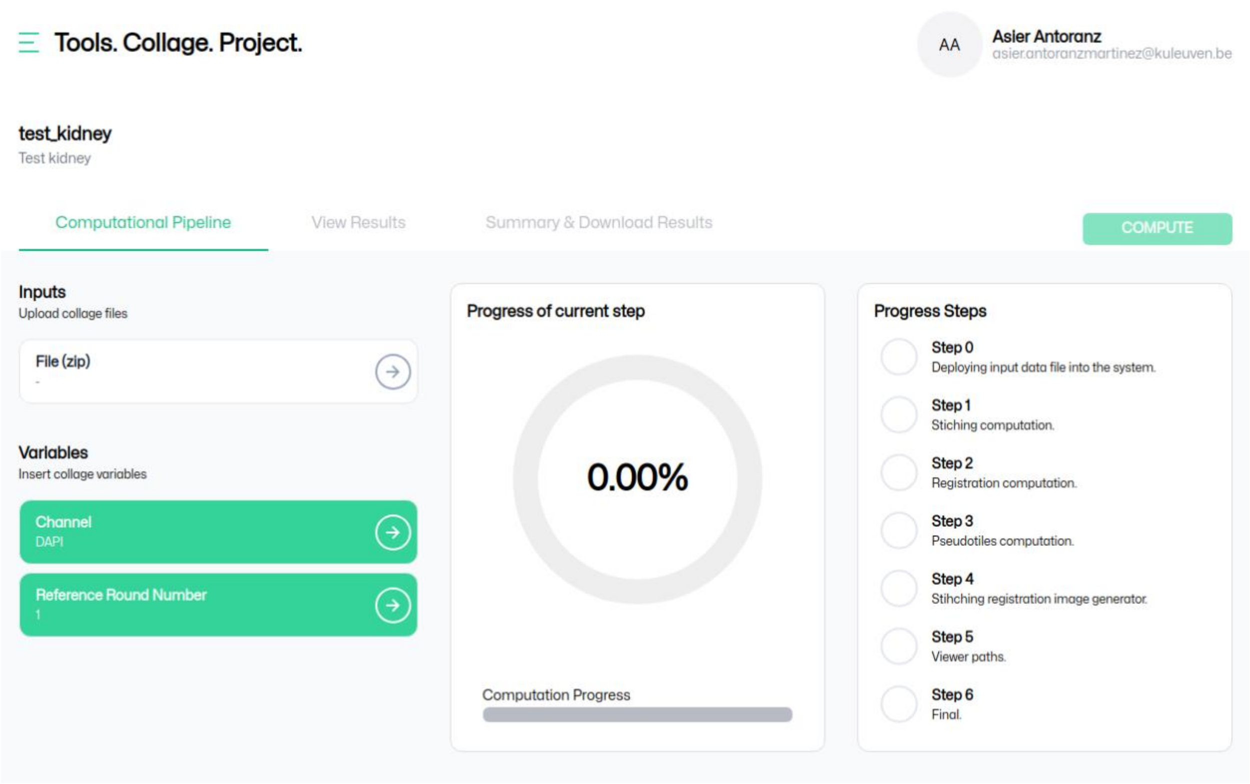


#### Data Upload

Next, the user should upload its data in .zip format. To illustrate the input data format, we will use some demo data we have deposited in Zenodo: https://zenodo.org/records/12698722. This .zip file has two folders corresponding to two cycles of a single sample. The folder nomenclature is as follows:

<slide_id\>_<cycle_id\>_ <version_id\>_ <project_id\>_ <user_id\>_ <tissue_id\>

For example:

KID1_R01_V1_KIDNEY_YVH_S0M

KID1_R02_V1_KIDNEY_YVH_S0M

Each of these folders contains a series of .tiff files corresponding to the acquired tiles in each channel and a csv file with the metadata corresponding to each tile. The tile nomenclature is as follows:

<slide_id\>_<cycle_id\>_ <version_id\>_ <project_id\>_ <user_id\>_ <mosaic_index\>_<channel_id/>.tiff

For example:

KID1_R01_V1_KIDNEY_YVH_S0M0_DAPI.tiff

The metadata file is a csv with the same name as the folder and the following mandatory attributes (note: more optional parameters are included but not needed):

- **C:** channel identifier in numeric format (0, 1, 2, etc.).
- **Frame:** It defines the size of the tile. For example, in this case, the tiles are squares of 2040 pixels. Therefore, the value is 4,4,2040,2040.
- **ImagePixelSize:** Pixel size, acquisition resolution for X and Y axes in 10*micrometers. In this case, the resolution is 0.65 microns. Therefore, the value in this field is ‘6.5,6.5’.
- **M:** tile identifier. It follows a numerical order starting from 0.
- **S:** tissue identifier. It follows a numerical order starting from 0.
- **StageXPosition:** Position of the stage across the X-axis in micrometers.
- **StageYPosition:** Position of the stage across the Y-axis in micrometers.
- **ValidBitsPerPixel:** Bit number of the images in numerical format. In this case, we have 16-bit images and therefore we write ‘16’.
- **Tile_filename:** Corresponding file name of the tile in the same path.

The upload can be done by clicking on the ‘File (zip)’ box below ‘Inputs’. This will make a window pop-up. You can drag and drop your .zip file and then click on ‘Upload’ to make the upload start. **Important note:** the upload won’t start until the user presses the ‘Upload’ button.

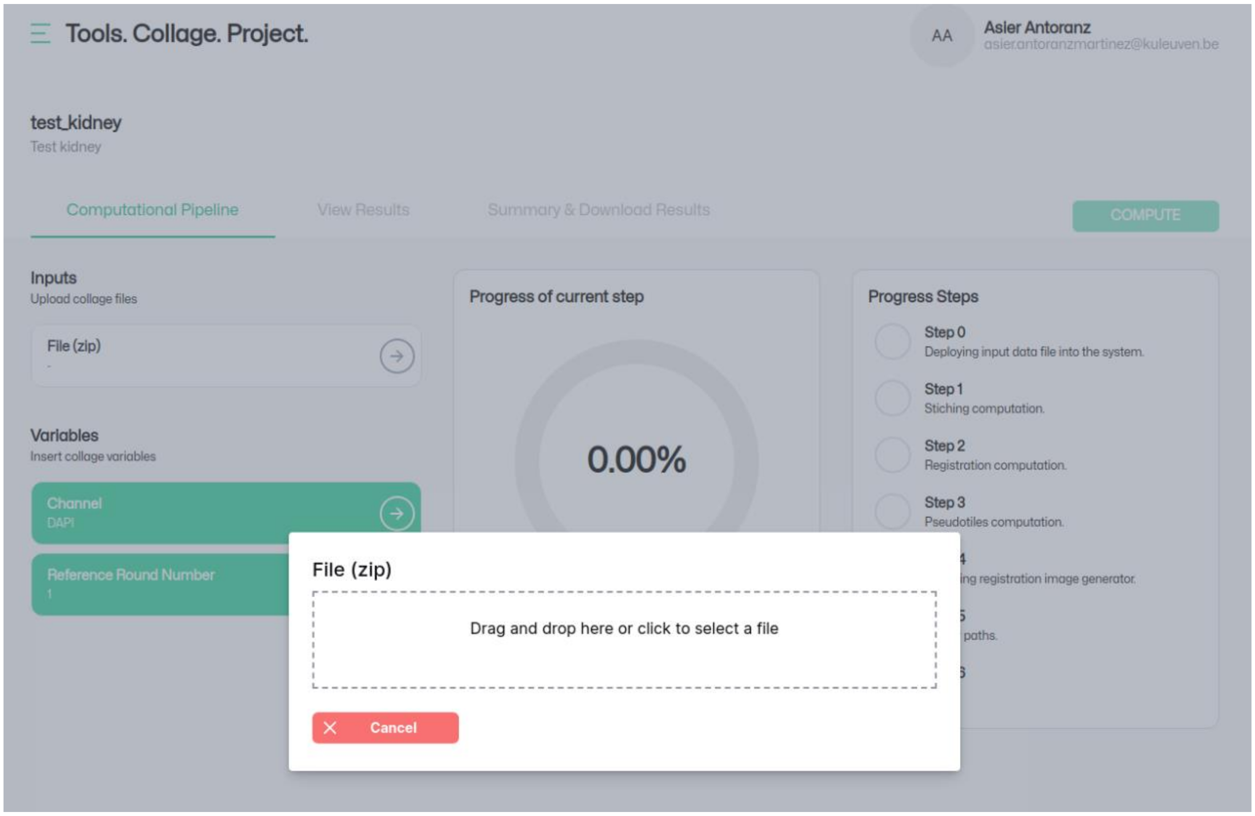


#### Parameter setting

After the file has been uploaded, we need to setup the parameters to run COLLAGE. There are two parameters:

- **Channel:** channel to be used for round alignment. We assume that different channels of the same staining round are aligned from acquisition. The standard value is DAPI. We recommend using DAPI or empty channels for the alignment. Staining channels obtain different markers each round and the different cycles might not be directly comparable hampering the registration.
- **Reference Round Number:** reference round to be used as fix image during Collage. The standard value should be the first round/cycle.

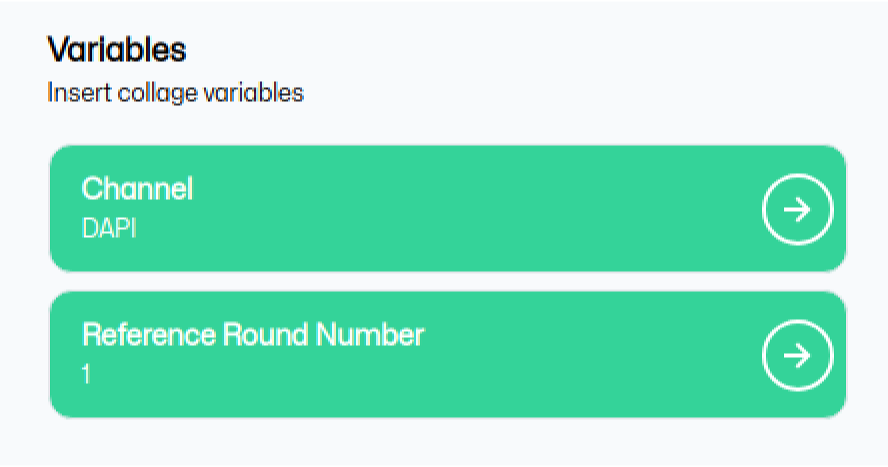


#### Analysis with Collage

In order to start with the analysis, press 'COMPUTE'. This will start COLLAGE. The user can follow the progress by checking the 'Progress Steps' section. The progress within each step is reported in the 'Progress of current step section'.

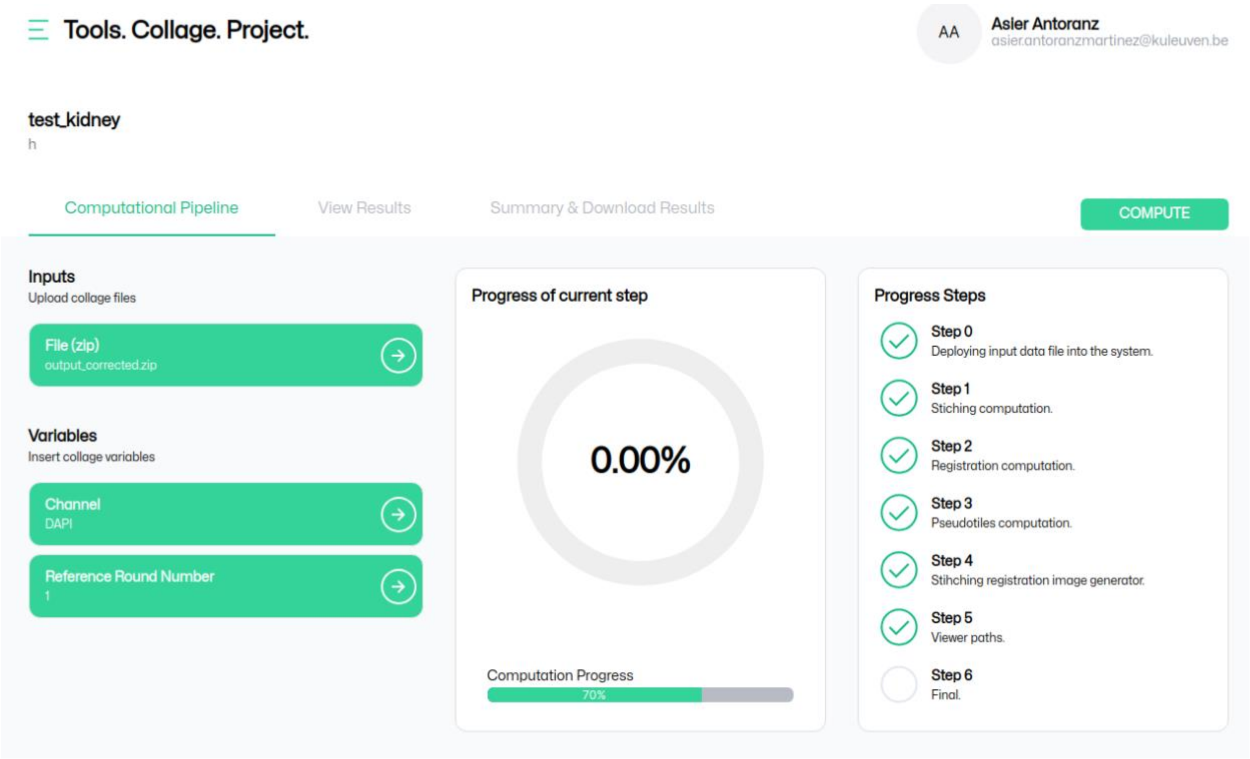


#### Visualization of Results

After the computation has finished, the user can access the 'View Results' section. There are three visualization modes:

- Composed_img: these are the stitched and registered images after Collage.

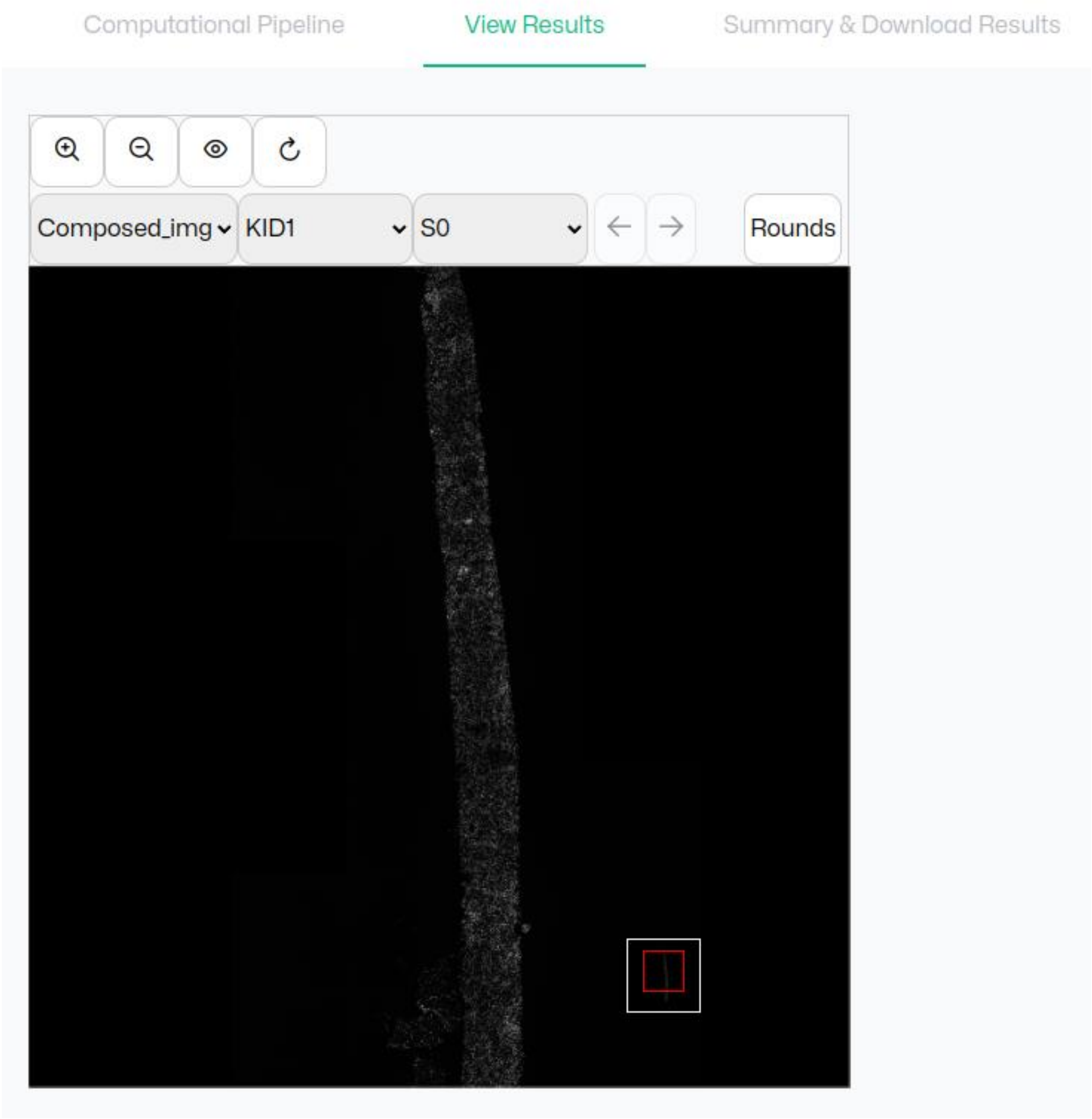


- Stitching_QC: these are quality control visualizations for the stitching. Stitching is only performed in the cycle selected in the parameter ‘Reference Round Number’.

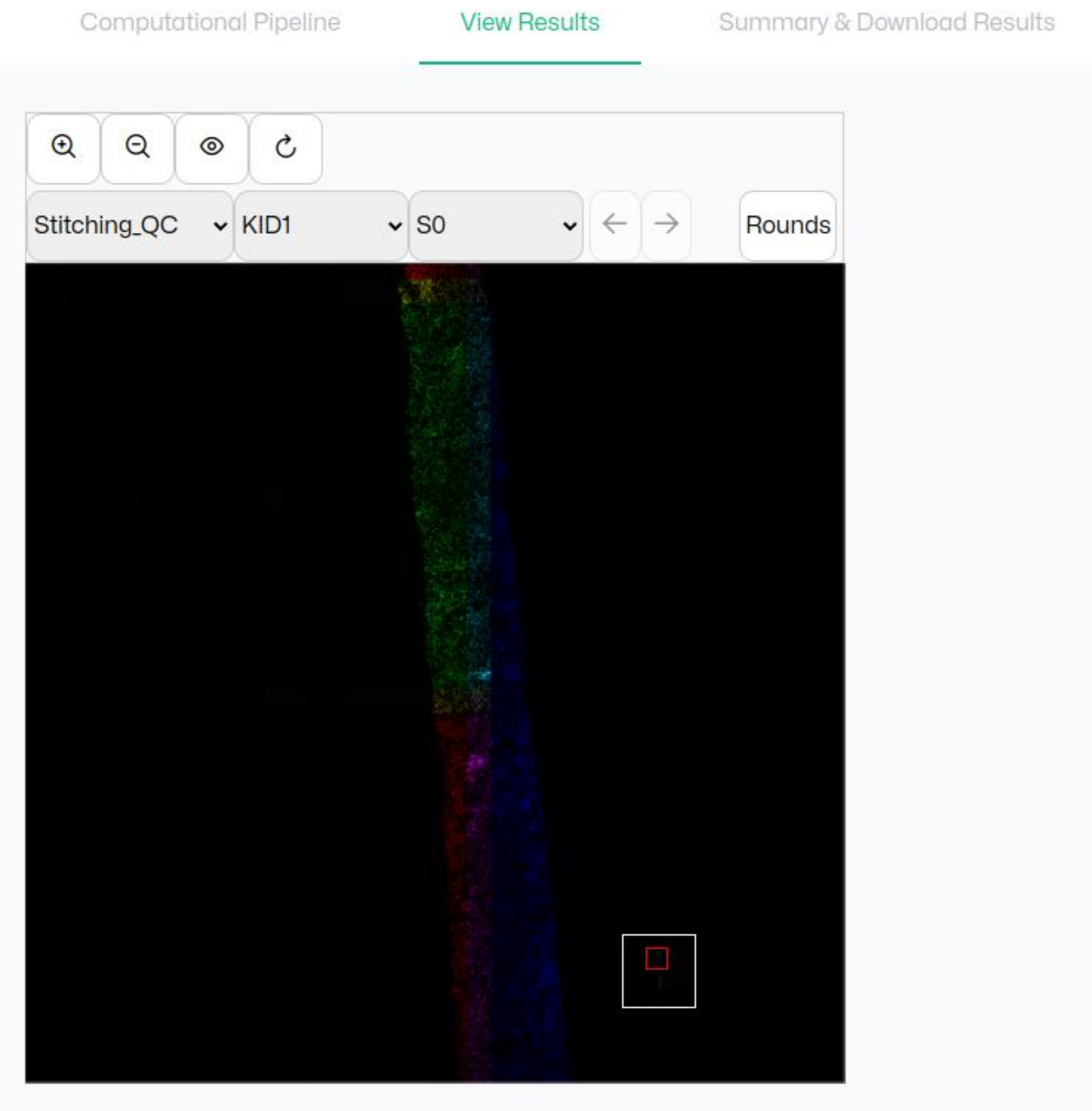


- Reg_Stitch_QC: these are quality control visualizations for the registration. These are generated for all rounds but only for the channel selected in the parameter ‘Channel’.

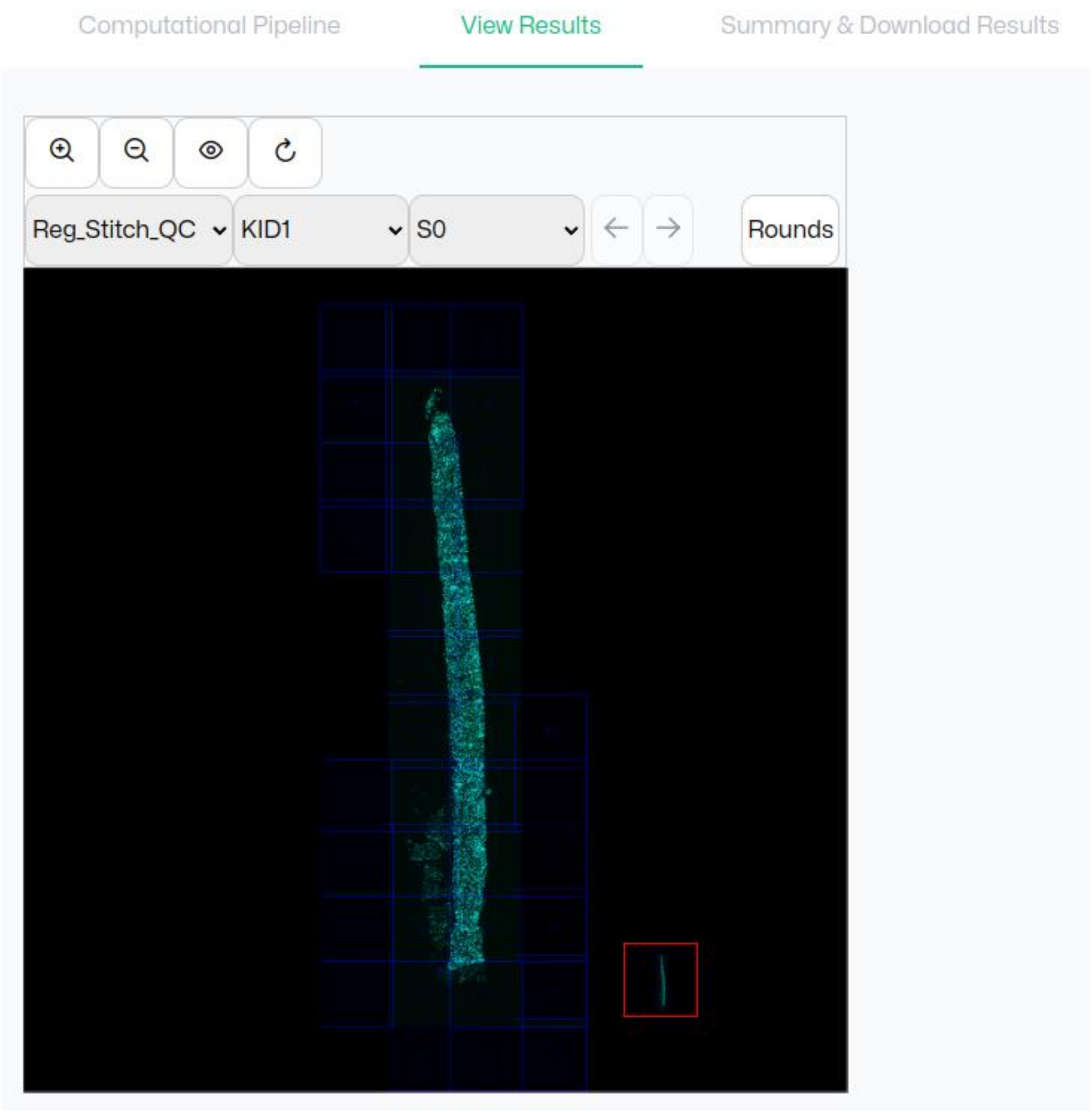


#### Download Results

The results obtained by Collage are downloadable via the ‘Summary & Download Results’ section. Files can be downloaded 1 by 1 or as a batch using the ‘DOWNLOAD ALL COMPOSED IMAGES’.

Composed_Img

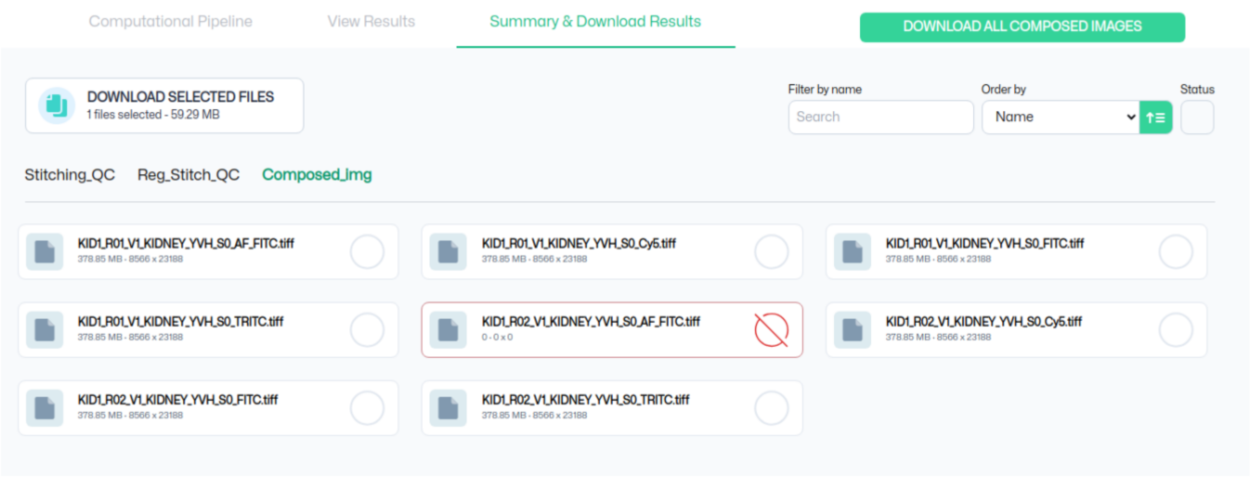


Stitching_QC

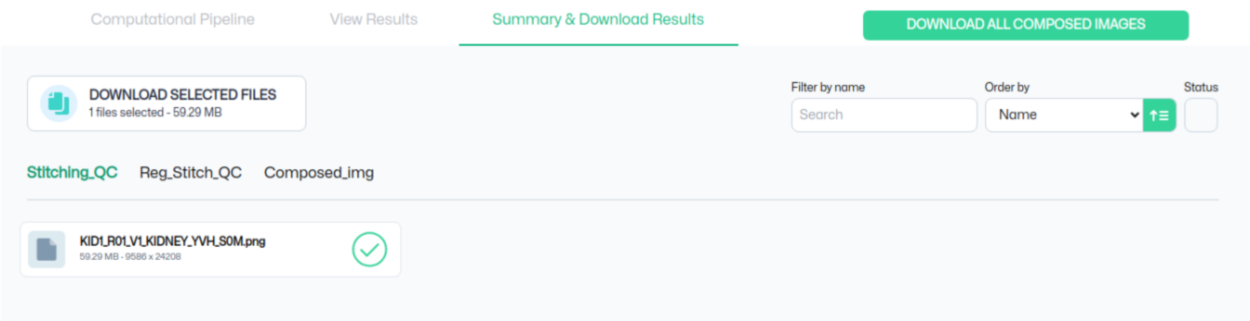


Reg_Stitch_QC

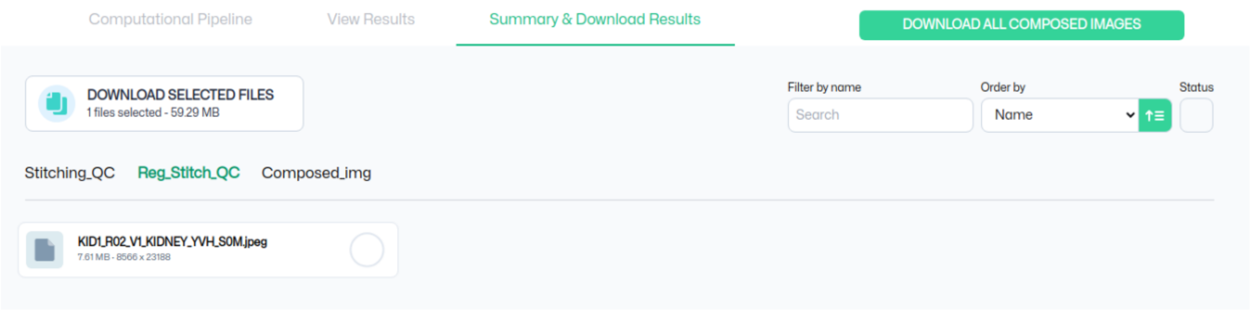
